# Representational dissimilarity component analysis (ReDisCA)

**DOI:** 10.1101/2024.02.01.578343

**Authors:** Alexei Ossadtchi, Ilia Semenkov, Anna Zhuravleva, Vladimir Kozunov, Oleg Serikov, Ekaterina Voloshina

**Author notes:** Corresponding author *Email address:* (Ekaterina Voloshina).

## Abstract

The principle of Representational Similarity Analysis (RSA) posits that neural representations reflect the structure of encoded information, allowing exploration of spatial and temporal organization of brain information processing. Traditional RSA when applied to EEG or MEG data faces challenges in accessing activation time series at the brain source level due to modeling complexities and insufficient geometric/anatomical data.

To address this, we introduce Representational Dissimilarity Component Analysis (ReDisCA), a method for estimating spatial-temporal components in EEG or MEG responses aligned with a target representational dissimilarity matrix (RDM). ReDisCA yields informative spatial filters and associated topographies, offering insights into the location of “representationally relevant” sources. Applied to evoked response time series, ReDisCA produces temporal source activation profiles with the desired RDM. Importantly, while ReDisCA does not require inverse modeling its output is consistent with EEG and MEG observation equation and can be used as an input to rigorous source localization procedures.

Demonstrating ReDisCA’s efficacy through simulations and comparison with conventional methods, we show superior source localization accuracy and apply the method to real EEG and MEG datasets, revealing physiologically plausible representational structures without inverse modeling. ReDisCA adds to the family of inverse modeling free methods such as independent component analysis [34], Spatial spectral decomposition [41], and Source power comodulation [9] designed for extraction sources with desired properties from EEG or MEG data. Extending its utility beyond EEG and MEG analysis, ReDisCA is likely to find application in fMRI data analysis and exploration of representational structures emerging in multilayered artificial neural networks.

## 1. Introduction

Representational Similarity Analysis (RSA) now stands as a pivotal technique within cognitive neuroscience, offering a profound lens into the organization and representation of information in the human brain. This method has revolutionized our capacity to comprehend neural bases of cognitive processes by elucidating relationships between neural representations of stimuli or concepts and relating them to behavioral variables and computational modeling [12, 30]. The largely philosophical concept of representational similarity later received its practical implementations in the field of both invasive electrophysiological [42, 19, 56] and non-invasive functional magnetic resonance imaging (fMRI) based neuroimaging [5, 8] followed by explicitly matching similarity patterns of neural activity to the theoretically pre-supposed representational structure [1] including the attempts to match the representational structure of neural activity across species [31]. Capitalizing on the success of the early application of the RSA principle, a seminal paper by Kriegskorte et al. [30] provided a step-by-step guide for the use of RSA to link the three whales of modern neuroscience: neuronal activity, behavior, and computational models. Since then a number of informative studies emerged that utilize the notion of representation [54] and employ RSA as an instrument to explore informational structure embodied in the neural substrate [29].

RSA was extended to merge fMRI and magnetoencephalography (MEG) non-invasive functional neuroimaging modalities [6]. This approach aimed to leverage the typically acknowledged high spatial resolution of fMRI alongside the high temporal resolution of MEG. MEG channels were used as features for constructing a classifier to yield the representational dissimilarity matrices (RDMs) corresponding to specific time intervals. These RDMs were then matched against those computed from spatial region of interest (ROI) activity patterns as measured by fMRI. This study revealed intriguing patterns of temporally persistent and recurrent representations [26] across the visual processing stream reflecting both bottom-up and top-down informational flows highlighting the concept of predictive coding implemented in our brains.

Several years later, MEG alone combined with cortical ROI multivariate pattern analysis and powered by the RSA framework [27] managed to provide both temporally and spatially informative maps highlighting the ventral vs. dorsal bifurcation of the visual pathway related to processing images of faces and tools contrasted against meaningless textures. Methodologically, this paper is significant as to the best of our knowledge that was the first attempt to apply RSA to MEG data in the source space. To this end, the authors first computed a fine-grained distributed inverse solution using the sLORETA technique [47]. Applying it to each time slice in the MEG sensor data they obtained cortical distribution of activity for each vertex on the cortical mesh model. Then, they combined vertices into spatially extended coarse ROIs and retained only three principal components of activities in each cortical ROI. Then, a series of region and time-window specific linear discriminant analysis (LDA) classifiers were trained to distinguish between stimulus classes and to form regional RDMs. These regional RDMs were then employed within the standard RSA framework and compared against theoretical RDMs.

It is important to realize that in the case of fMRI, the source space signals are readily available after the application of a more or less standard collection of analysis steps including data preprocessing and calculation of the appropriate contrasts. Things are significantly less transparent and standardized in the EEG or MEG cases [39, 10]. Here, to obtain the individual voxel or ROI activity profiles, the inverse modeling approaches are used to convert sensor signals into the source space activation time series. This is typically done by applying an inverse operator matrix to EEG / MEG sensor measurements. The inverse operator directly depends on the forward model. The latter approximates the way the source activity is mapped to the sensors and is computed by solving Maxwell equations within the head volume which typically requires approximating the head with a set of nested surfaces bounding the regions of constant conductivity. The accuracy of the forward model depends on the extent to which the geometry as well as conductivity and permeability properties of the model align with their real counterparts. Generally unknown conductivity profiles of the tissues comprising the head, the associated anisotropy of the conductivity tensor, and the limited coregistration accuracy of the sensor array to the volume conductor model are the sources of the forward model inaccuracies especially pronounced when dealing with EEG data. The precision of modeling magnetic fields from neuronal sources tends to yield accurate MEG forward models, showcasing an impressive fidelity with an estimated error rate of around 10% [38]. Yet, despite this advantage, MEG systems are notably less accessible compared to their EEG counterparts. Even if the forward model is accurate enough the task of estimating neuronal sources from non-invasively collected EEG or MEG data is inherently ill-posed. Fundamentally, for a given set of measurements, there exist an infinite number of cortical source distributions that perfectly fit the measurements. This problem is resolved and a unique solution is guaranteed by regularization which makes the results depend heavily on the *apriori* assumptions about source distributions. Taken together, all this limits RSA applications in the realm of time-resolved neuroimaging furnished by the EEG and MEG, the techniques that offer a unique time-resolved window into the rapid neuronal processes.

To address this problem and broaden the potential applications of the powerful RSA principle here we propose novel representational dissimilarity component analysis (ReDisCA) - an inverse modeling free method designed to estimate the spatial-temporal components in evoked responses that adhere to a specific target (or theoretical) representational dissimilarity matrix (RDM). The output of this method is a set of spatial filters and the associated topographies (or patterns) that appear informative of the location of “representationally relevant” sources whose activity exhibits the sought representational profiles. Application of the discovered spatial filters to the evoked response time series matrix yields temporal source activation profiles with the desired RDM over the selected time window. It is worth noting that ReDisCA belongs to the class of spatial decomposition methods that resolve the inherent uncertainty of the inverse problem by adding the specific assumptions and looking for the number of sources or components whose count does not exceed the rank of the data matrix. Thus, ReDisCA looks for components (or sources) with specific representational dissimilarity profiles.

While ReDisCA does not necessitate an inverse modeling, its foundation lies in the classical linear EEG/MEG observation equation. It exercises the concept of spatial filtering, akin to already established techniques like Independent Component Analysis (ICA) [34], Spatial-Spectral Decomposition (SSD) [41], or Source Power Co-modulation (SPoC) [9]. This ensures that the discovered spatial patterns are suitable for implementing a rigorous source localization procedure based on the notion of signal subspace [40] as described for example in [44] in application to ICA.

This compatibility with the linear model positively distinguishes ReDisCA from the purely sensor space applications of RSA to EEG or MEG data. For example, [37] used high-density EEG and highlighted the detailed spatial and temporal patterns of responses underlying various aspects of meaningfulness in the presented visual stimuli. They achieved this by using newly proposed differentiation analysis (DA) and computed temporal and spatial profiles of the category differentiation index (CDI) that can be used to make judgments about time windows and approximate spatial locations of the pivotal sources. However, due to their empirical nature, the obtained spatial CDI maps can not be rigorously fitted with the electromagnetic model of the head and therefore are only tangentially related to the underlying distribution of cortical sources with target representational properties. Therefore, the approach proposed here can be considered as a link between the existing purely sensor-space applications of RSA to EEG and MEG data and the fully-fledged inverse modeling-based technique described in [27].

In what follows we first describe a classical RSA formulation and then proceed to the presentation of ReDisCA. As it will become clear, ReDisCA utilizes formally the same optimization strategy as that used in SPoC, a well-established technique for extracting spatial-temporal components corresponding to neuronal sources of rhythmic activity whose power is co-modulated with the user-supplied behavioral variable. To illustrate this connection we provide a table of explicit correspondence between the quantities defined in this paper and those used in SPoC and then describe the optimization strategy based on solving the generalized eigenvalue problem with appropriately formed matrices. Next, we evaluate ReDisCA’s performance and compare it against several versions of the source space RSA implementations in the two simulated scenarios, with either a single source or with four simultaneously active sources each with its own RDM. Finally, we apply ReDisCA to the analysis of a real EEG dataset that lacks the information necessary for source localization, a prerequisite for the source space RSA analysis. We show how the new method can be used to discover representationally relevant structure in the EEG and MEG evoked responses without the use of inverse modeling. We conclude with a discussion of ReDisCA’s strengths and weaknesses and relate it to the existing modifications of RSA including accommodation of advanced dissimilarity measures [58] and the data-driven estimation of the theoretical RDM [25, 21].

## 2. Methods

### 2.1. Problem statement

Consider a collection of data matrices denoted as 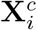, each with dimensions *N* × *T*. These matrices represent recordings of brain activity, specifically EEG, MEG, or potentially ECoG or sEEG, performed with *N* channels over *T* time stamps. The length of the time segment of interest within the entire response is *T* samples, corresponding to the number of columns in 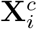. These recordings are obtained during the *i*-th trial out of a total of *I*_*c*_ trials, corresponding to the experimental condition *c* = 1, …, *C*.

Averaged evoked response is one of the informative features extracted from electrophysiological recordings obtained in the event-related paradigms. It can be calculated simply by averaging 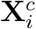 over *I*_*c*_ trials to get 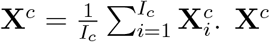 emphasizes phasic components of the response, i.e. those whose activity phase is consistent with the stimulus onset.

Our objective is to use a set of 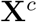 observed over *C* conditions and to identify spatial and temporal components or subsequently brain regions whose temporal activity profiles exhibit the target differences among the *C* experimental conditions. The target differences in our case are defined by a predefined theoretical or model-based *C* × *C* representational dissimilarity matrix (RDM) 𝔻.

### 2.2. Source space representational similarity analysis (RSA)

For EEG and MEG scenarios, we can achieve this by the two-step spot-light procedure similar to that used in [27]. First we use an *M* × *N* inverse operator denoted as **W**_*M*_ whose *m*-th row is denoted as 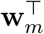, *m* ∈ 1, …, *M*, and acts as a spatial filter tuned to the *m*-th voxel or cortical vertex.

This precomputed spatial filter aids in estimating the 1 × *T* vector of the *m*-th voxel activity during condition *c* as 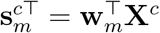, allowing us to derive the time series corresponding to every voxel in the predefined volumetric grid based on the sensor-space averaged evoked response measurements **X**^*c*^ defined above. Note that here and in the subsequent expressions horizontally oriented vectors are marked with transpose superscript, i.e.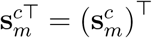.

Next, once having obtained regional time series vectors 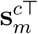 we can evaluate the dissimilarity measure of choice to obtain the *m*-th region-specific dissimilarity matrix 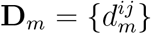, *m* = 1, …, *M*. In what follows we will stick to the square of the Euclidean distance as a measure of response dissimilarity that can be readily extended to Mahalanobis distance once the covariance matrix describing the dependencies between the time samples of the regional response is available. Thus

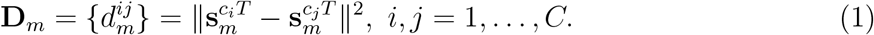

Once the regional RDMs are prepared, following the standard RSA procedure, we will employ the correlation coefficient *ρ*_*m*_ to assess the similarity between regional RDMs (**D**_*m*_) indexed by *m* and the theoretical RDM 𝔻 = *{d*^*ij*^*}* typically supplied by the user.

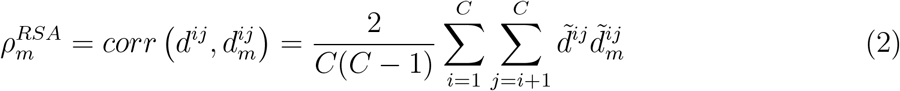

where the tilde represents standardized upper triangular elements of the respective RDM matrices. The standardization operation applied to a vector lies in subtracting its mean and dividing by the standard deviation.

The scores 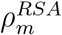 reflect the similarity between the theoretical and regional RDMs. They can be visually represented on the cortical surface or within the brain volume by interpolating the scores associated with each of the *M* voxels or cortical mesh nodes. Then, to identify the spots of significant similarity, a statistical testing procedure is applied. This procedure should consider the limited spatial resolution inherent in EEG and MEG-based inverse mapping, and correction should be made for the effective number of multiple comparisons. This may be accomplished using, for example, the cluster-based permutation testing procedure [36] where the surrogate data are obtained by permuting the labels of *C* conditions which essentially corresponds to computing 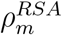 but for a randomly reshuffled upper triangle of the theoretical RDM.

The diagrams in Figure 1 schematically show the described process. In the first case illustrated in panel Figure 1.a an inverse operator is applied to already averaged evoked response data **X**^*p*^ observed during condition *p, p* = 1, …, *C*, and then the empirical RDM entry is calculated as the quadratic difference between the evoked source time series in condition *p* and *q*. In Figure 1.b we show an alternative approach where the inverse operator is applied to the single (*l* − *th*) trial data, followed by computing the quadratic difference that then gets averaged over trials within each condition. In our experiments with source space RSA, we will also distinguish between the use of two different inverse solvers - Minimum norm estimator (MNE) and Linearly constrained minimum variance beamformer (LCMV BF). This gives us in total four versions of the source space RSA that we will compare our new approach with.

**Figure 1:**
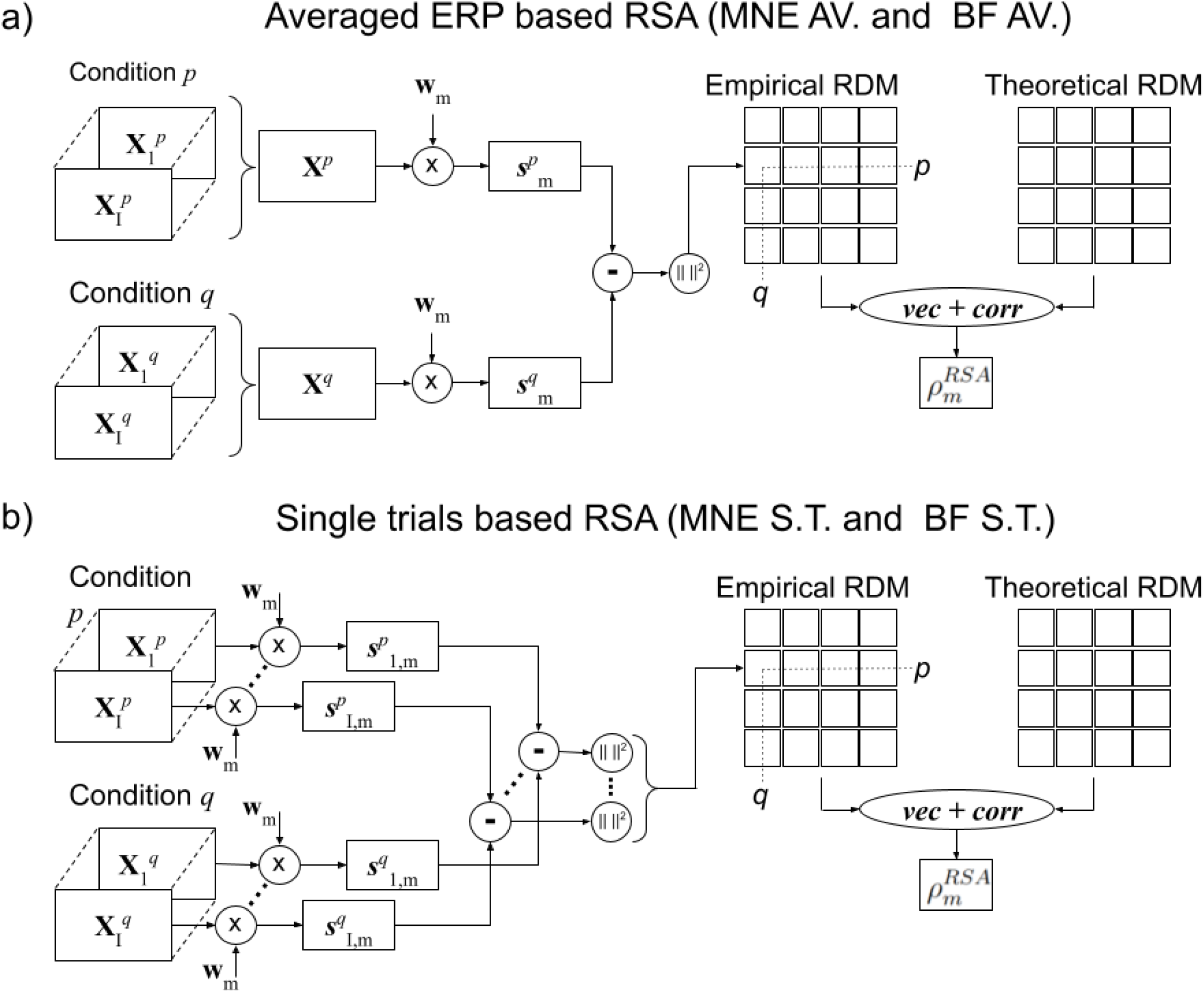
Source space RSA diagrams. a) RSA applied to the averaged evoked response (AV). The inverse operator is applied to already averaged evoked response data **X**^*p*^ observed during condition *p, p* = 1, …, *C*, then the empirical RDM entry is calculated as the quadratic difference between the evoked source time series in conditions *c*_*i*_ = *p* and *c*_*j*_ = *q*. b) RSA applied to single trials (S.T.). The inverse operator is applied to every single *l* − *th* trial in the data, followed by computing the quadratic difference that gets then averaged over trials within each condition. In both cases, different inverse operators 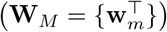 can be used. We experimented with two inverse solvers - MNE and LCMV BF.

### 2.3. Representational dissimilarity component analysis (ReDisCA)

Here we propose an alternative method for identifying brain activity components with the desired representational dissimilarity. Instead of using the pre-computed inverse operator 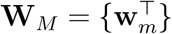, *m* = 1, …, *M* whose *m*-th row serves the *m*-th cortical vertex and conducting an exhaustive search over all *M* cortical vertices (or voxels) as in the spotlight source space RSA illustrated in Figure 1, we analytically seek a vector of coefficients **w** such that the empirical RDM of spatially filtered data 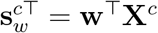, *c* ∈ 1, …, *C*, closely approximates the target RDM 𝔻. Note that this analytically found spatial filter weights vector **w** conceptually plays the same role as the rows **w**_*m*_ of the inverse operator in the source space RSA.

To implement this we posit and solve the following optimization problem

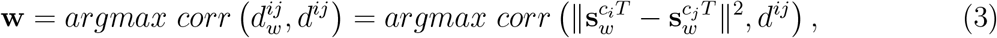

where following (1) to gauge similarity between the empirical RDM 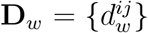 of the spatially filtered data 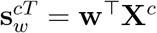 and the target RDM matrix 𝔻 we use the Pearson correlation coefficient between the two RDMs upper triangular elements indexed by (*i, j*). Also, as a representational dissimilarity measure we employ the squared Euclidean distance between condition-specific spatially filtered temporal activation profiles. However, in this case, the spatial filter **w** is unknown and needs to be found by solving this optimization problem. Once the weights are found they can be turned into source topographies. The relation between spatial filter weights and the associated source topographies has been described in [17] and will also be addressed below. The obtained topographies of sources with desired RDM profiles can then be used for inverse modeling and localization of the underlying neuronal sources. Note that we used subscript *w* to denote the elements of the empirical RDM corresponding to the spatial filter **w** that we are about to identify, see (2) for comparison.

To solve the optimization problem (3) we use the fact that 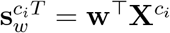 and rewrite (3) as an explicit function of **w**:

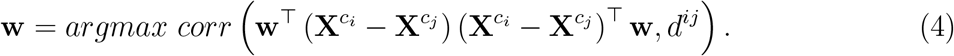

For compactness, we denote 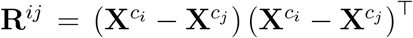 to obtain the following optimization problem

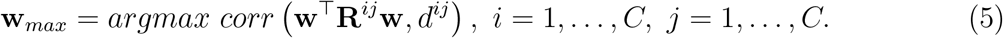

In Equation (5), *N* × *N* matrix **R**^*ij*^ represents the unscaled correlation matrix of the sensor-space time series differences recorded between condition *i* and condition *j*. This expression formally exactly matches the optimization problem solved in SPoC [9] where the role of **R**^*ij*^ is played by the time-segment specific sensor data correlation matrix.

SPoC deals with a collection of time segments (or epochs) indexed by *e* in the original paper, and aims to find a spatial filter for isolating a source whose power follows the desired temporal profile *z*(*e*). In our approach, we work with a collection of condition pairs (*i, j*) and seek a source whose RDM 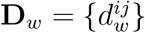 matches the target RDM 𝔻 = {*d*^*ij*^}. While SPoC computes correlation matrices **C**(*e*) specific to the *e*-th epoch of EEG(MEG) data, in our approach we calculate correlation matrices **R**^*ij*^ of the difference of EEG (MEG) time series observed during the pairs of conditions indexed by (*i, j*). The role of index *e* in the original SPoC is played by the pair of indices (*i, j*) whose combinations enumerate the collection of the upper triangular elements of the RDM. One may also express this correspondence as *e* = (*i* − 1)*C* + *j, i > j, i* = 1, …, *C, j* = 1, …, *C*. Essentially, the *e*-th segment time series in SPoC is replaced by the time series of the difference between condition pair (*i, j*). The role of the desired power profile *z*(*e*) from the original SPoC is played by the elements *d*^*ij*^ of the theoretical RDM 𝔻 driving the proposed here representational dissimilarity components analysis (ReDisCA). This correspondence is summarized in Table 1 below.

**Table 1:** Correspondence between main SPoC variables as defined in [9] and the variables used to define ReDisCA in this manuscript.

| Var. SPoC | Var. ReDisCA | Role in SPoC | Role in ReDisCA |
| --- | --- | --- | --- |
| $e$ | $(i, j)$ | time window index | conditions pair index |
| $\mathbf{C}(e)$ | $\mathbf{R}^{ij}$ | data correlation matrix | data diff. corr. matrix |
| $z(e)$ | $d^{ij}$ | desired response pattern | desired RDM pattern |

There are two ways to solve the optimization problem (5). The first one is to use the gradient descent and to identify **w** as the solution for (5) to maximize the correlation coefficient between the empirical and the desired theoretical RDM profiles. While this is entirely possible, the SPoC paper describes a more attractive alternative to approximate the correlation coefficient with covariance which allows for a closed-form solution.

To ensure that optimization of covariance (the non-normalized version of the correlation coefficient) is a good approximator of the correlation coefficient optimization we

- use standardized elements of the RDM 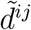
- use the constraint 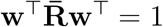 where

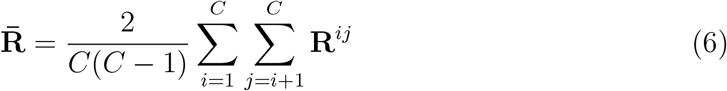

is the average correlation matrix.

For compactness we also denote the weighted average correlation matrix

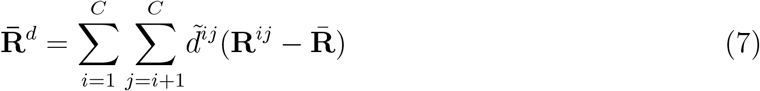

The resulting optimization problem can then be written as

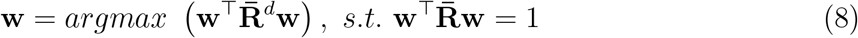

and can be analytically solved using the method of Lagrange multipliers.

The problems formally identical to (8) often arise in the domain of multivariate signal processing, see beamforming [57], common spatial patterns [49], spatial spectral decomposition (SSD) [41] and SPoC [9] as EEG (MEG) specific examples. The optimal **w** is found as the solution to the following generalized eigenvalue problem corresponding to the largest eigenvalue

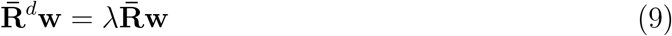

When solving this problem for an *N* -channel data we will arrive at an *N* ×*N* matrix **W** whose columns are the eigenvectors of matrix pair 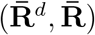 or in other words

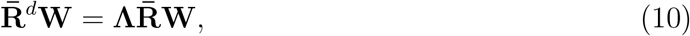

where **Λ** = {*λ*_*n*_ }, *n* = 1, …, *N* is the diagonal matrix of generalized eigenvalues. The above procedure is referred to as the maximization of covariance option in the original SPoC paper [9]. In this case, matrix **W** resulting from solving the generalized eigenvalue problem (10) is square and also full rank as we assume that both 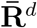 and 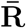 are invertible. Then, **W** can be simply inverted to obtain a matrix 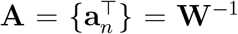 whose rows 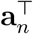 are the topographies of sources with the desired representational properties. In the case when the generalized eigenvalue problem (10) is not full rank the described procedure can be performed in the lower dimensional principal space and the obtained topographies can be transformed back to the original sensor space.

Note, that by solving (10) instead of (9) we obtain *N* spatial filters **w**_*n*_ and source topographies 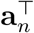, *n* = 1, …, *N*. The *n*-th generalized eigenvalue *λ*_*n*_, *n* = 1, …, *N* reflects the extent to which the empirical RDM corresponding to the time series extracted using spatial filter **w**_*n*_ is similar to the desired theoretical RDM 𝔻.

We call the topographies 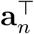 combined with the corresponding data derived RDM profiles 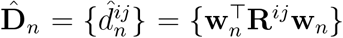 the *representational dissimilarity components* and the described process of obtaining them the *REpresentational DISsimilarity Component Analysis* or ReDisCA.

The significance of the similarity between the component-specific data-derived RDMs 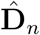, *n* = 1, …, *N* and the theoretical RDM 𝔻 can be established by the permutation testing procedure suggested in SPoC. The idea of this approach is to solve a large number of surrogate problems of the form (10) but based on the matrices created from the data with permuted condition labels. The mutual correspondence between the set of difference correlation matrices **R**^*ij*^ and the condition pair labels (*i, j*) is destroyed and the corresponding asymptotic *p*− values are calculated as a fraction of cases when the surrogate generalized eigenvalue exceeds that obtained on the original data.

The collection of *K* topographies **A**_*K*_ = [**a**_1_, …, **a**_*K*_] corresponding to the components with significantly high similarity between the data-derived and the theoretical RDM spans *representational dissimilarity subspace, ReDisS*. These topographies together with the corresponding spatially filtered ERP time series furnish the decomposition of the evoked response potential (ERP) data into a set of *relevant* constituents with respect to the used theoretical RDM.

Technically this completes ReDisCA’s description whose result is a set of spatial component topographies **a**_*k*_ and the corresponding spatial filters **w**_*k*_ that can be used to obtain component-specific response time series vectors 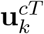 in the *c*-th condition as 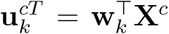. More generally, one can write this in a matrix form as

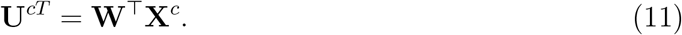

From the above and given that the matrix of spatial filters **W**^⊤^ is invertible we can write

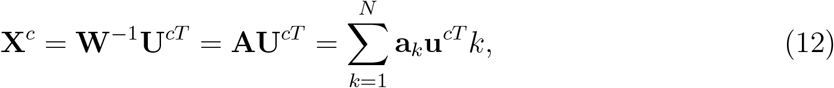

which shows that after obtaining ReDisCA components the sensor data in the *c*-th condition can be reconstructed as a superposition of rank-1 contributions **a**_*k*_**u**_*kc*_ of ReDisCA components.

Note that using ReDisCA does not require an explicit electromagnetically informed inverse modeling to identify the spatial components within multichannel EEG (MEG) data that exhibit the desired representational properties aligned with the user-supplied theoretical RDM.

However, to compare ReDisCA’s results to those obtained with the more conventional source space RSA approaches we can apply the electromagnetic inverse modeling to the identified spatial topography vectors **a**_*k*_. This allows us to quantitatively assess the locations of sources underlying each of the components. The simplest way to associate ReDisCA results with neuronal sources is to compute the cosine similarity score between each forward model-based topography vector **g**_*m*_, *m* = 1, …, *M* corresponding to the *m*−th vertex of the cortical mesh and the ReDisCA derived pattern **a**_1_ corresponding to the largest generalized eigenvalue *λ*_1_, i.e.

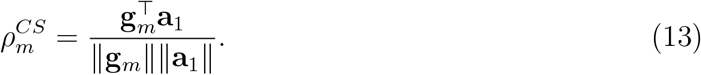

This procedure can be applied to each statistically significant topography.

Alternatively, to localize sources corresponding to the entire *K* − dimensional representational dissimilarity subspace spanned by the topographies of statistically significant components stored in **A**_*K*_ we can perform a simple multiple signal classification (MUSIC) scan using subspace correlation metric as

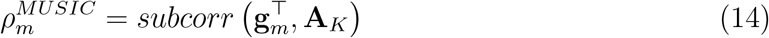

Similarly to *ρ*_*m*_ used in the classical RSA setting, the scores 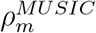 and 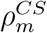 can be visually represented on the cortical surface or within the brain volume by interpolating the scores associated with each of the *M* voxels or cortical mesh nodes. Note also that 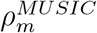 is a more general measure of similarity between the subspaces of the arbitrary dimensions and reduces to 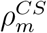 for *K* = 1.

### 2.4. Simulations setting

To validate the proposed ReDisCA approach, we conducted two sets of simulations. The best way to ensure that ReDisCA derived components are informative with respect to the underlying sources is to actually find these sources given the components estimated by ReDisCA. Therefore in both simulations we mapped the obtained topographies to the source space using (14) and assessed the location of the MUSIC scan’s peak with respect to the simulated source location(s).

#### 2.4.1. Generation of source activation time series with desired representational structure

In both simulations we used the following strategy to generate the source activation time series for each of the *C* conditions and associate them with the representational structure across *C* conditions.

- For each source generate a random mixing matrix **M** by sampling its elements from zero mean and unit variance Gaussian distribution 𝒩 (0, 1)
- Generate *C* × *T* matrix **S** of condition-specific source evoked response potentials as **S** = **MZ** where **Z** is a *C* ×*T* matrix of Gaussian random variables whose rows are low-pass filtered by the 6-th order Butterworth filter with a cut-off frequency of 2 Hz. We will refer to the *c*-th row of this matrix as 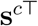 for *c* = 1, …, *C*.
- Compute the resultant dissimilarity profile matrix 𝔻^0^ from the obtained condition-specific time series vectors **S** according to equation (1).

As described later the noisy version of 𝔻^0^ will be supplied to the tested here RSA approaches as a theoretical RDM. For each Monte-Carlo iteration we generated a new matrix **Z**, and thus a new set of condition-specific source time series **S**. In the first set of simulations where we generated only one active source we kept the mixing matrix *M* constant over all Monte-Carlo iterations. In the second set of simulations with *P* = 4 sources, we used four mixing matrices **M**_*p*_, one for each source, that remained unchanged throughout Monte-Carlo trials.

#### 2.4.2. ReDisCA vs. source space RSA

Within the first set of simulations we compared ReDisCA against the source space RSA using a single source scenario. In this comparison, to generate the observed EEG (MEG) data we placed a single source iteratively in a node of the cortical mesh whose index was randomly generated and activated the source with the condition-specific time series. Here we used *C* = 5 conditions and generated condition-specific ERP time series. For each source position (iteration) the time series were randomly regenerated from a newly sampled matrix of random variables according to the procedure described above.

At each Monte-Carlo iteration corresponding to a specific source location in the *m*_0_-th node of the cortical mesh we obtained condition-specific source time series observed in the *l*−th trial as

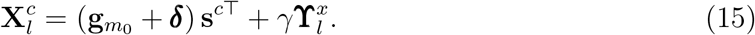

In (15) 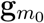 is the *N* × 1 forward model vector (or topography) corresponding to the source located in the *m*_0_-th vertex of the cortical mesh with coordinate 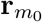. In order to account for inevitable forward modeling errors we used normally distributed *N* × 1 source model noise vector ***δ*** with covariance matrix 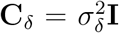 and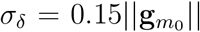. Row vector 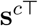 is the *c*-th row of **S** and 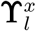 is a *N* × *T* realistic brain noise matrix generated by 1000 randomly seeded cortical sources activated with 1*/f* noise, similar to the way it was done in [43] with factor *γ* controlling the signal-to-noise ratio in the simulated data. Note that each trial had a different realization of the spatially correlated noise 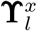 and new ***δ*** was generated for each Monte-Carlo iteration.

We then use the simulated noisy sensor data 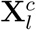 for *l* = 1, …, *I*_*c*_ and *c* = 1, …, *C* and we aim to use the approximate RDM of a specific source to find spatial component(s) whose time series, see (11), exhibits the desired representational structure defined by the approximate RDM. The approximate RDM is used to simulate the real-world scenario where we naturally lack precise knowledge of the dissimilarity profile. To this end, we incorporate a random *C* × *C* noise matrix **ϒ**^*d*^ at each Monte-Carlo iteration. This addition modifies the theoretical RDM 𝔻^0^ to 𝔻 = 𝔻^0^ + **ϒ**^*d*^.

To verify the result we then apply the source localization procedure to the discovered component(s) to determine the location 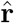 of a source (or its index, denoted as 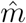 the discovered spatial component pertains to and whose activity exhibits the intended theoretical similarity profile. We then use the obtained location to calculate the performance metrics as described in Section 3.1. When source space RSA is applied to this simulated data our goal is to directly identify the cortical source of interest using the procedures outlined in Figure 1.

In our simulations we have chosen *T* = 200 ms as the length of the window to which RSA is applied. The spectral profiles of the source evoked activity occupied the frequency range below 2 Hz to impose the smoothness characteristic of the real evoked responses. As we will show in the example of applying ReDisCA to real EEG data the method can be applied in a sliding window mode and remains operable at *T* = 150 ms. The constraint on the length of the time interval is dictated primarily by the condition number of the resultant correlation matrices (4) and can be further reduced with proper regularization and/or dimension reduction procedures. Alternatively, ReDisCA can be applied to the entire evoked response time interval and then a statistical testing procedure can be used to determine the intervals of significant difference, see Section 4.2.2.

To evaluate the performance of ReDisCA in comparison to the four versions of the source space RSA, we employed the Receiver Operating Characteristic (ROC) curve. This measure, devoid of specific thresholds, gauges the overall informational capacity in the output of these approaches concerning the detection task. For a detailed explanation of how the ROC curve is calculated, please refer to Section 3.1.

#### 2.4.3. Realistic simulations with multiple sources

Another simulation scenario reproduces the situation when the observed evoked responses are generated by a superposition of several sources each with its own representational dissimilarity matrix. To this end we experimented with *P* = 4 sources and their corresponding dissimilarity matrices 𝔻^0*p*^ and their noisy versions 𝔻^*p*^, *p* = 1, …, *P*. The ideal RDMs 𝔻^0*p*^, *p* = 1, …, *P* of the evoked responses for each of *P* = 4 sources among *C* = 6 conditions are shown in Figure 3. To evaluate the RSA’s robustness we have also experimented with a subset of *C* = 3, 4, 5 conditions.

At each Monte-Carlo iteration we have randomly seeded four sources over the cortical mantle with the restriction of no source being closer than *δ*_*min*_ = 2 *cm* to any of the other sources. This gave us *P* = 4 true source location vectors 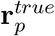 and true source topography vectors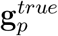, *p* = 1, …, *P*. We have then added noise to these true source topographies, projected source activation time series to the sensors and added realistic 1*/f* brain noise similarly to (15) as

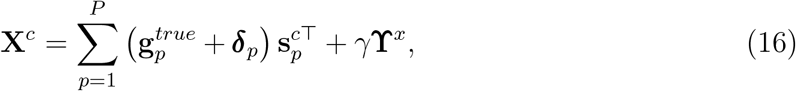

Overall the approach to constructing the simulated data in this second experiment matched that of the first one with the exception that in the current experiment the simulated data contained the superposition of activity of *P* = 4 simultaneously active task-related sources each with its own RDM.

## 3. Performance metrics

As we described in Section 2 ReDisCA itself does not require the electromagnetic inverse modeling, see also Figure 2. At the same time the source space RSA applied to EEG(MEG) data explicitly uses inverse modeling as highlighted by the diagrams shown in Figure 1. Therefore, to evaluate ReDisCA’s performance and match it against the source space RSA we use components discovered by ReDisCA and fit them with the electromagnetic model as described in Section 2 and equations (13, 14).

**Figure 2:**
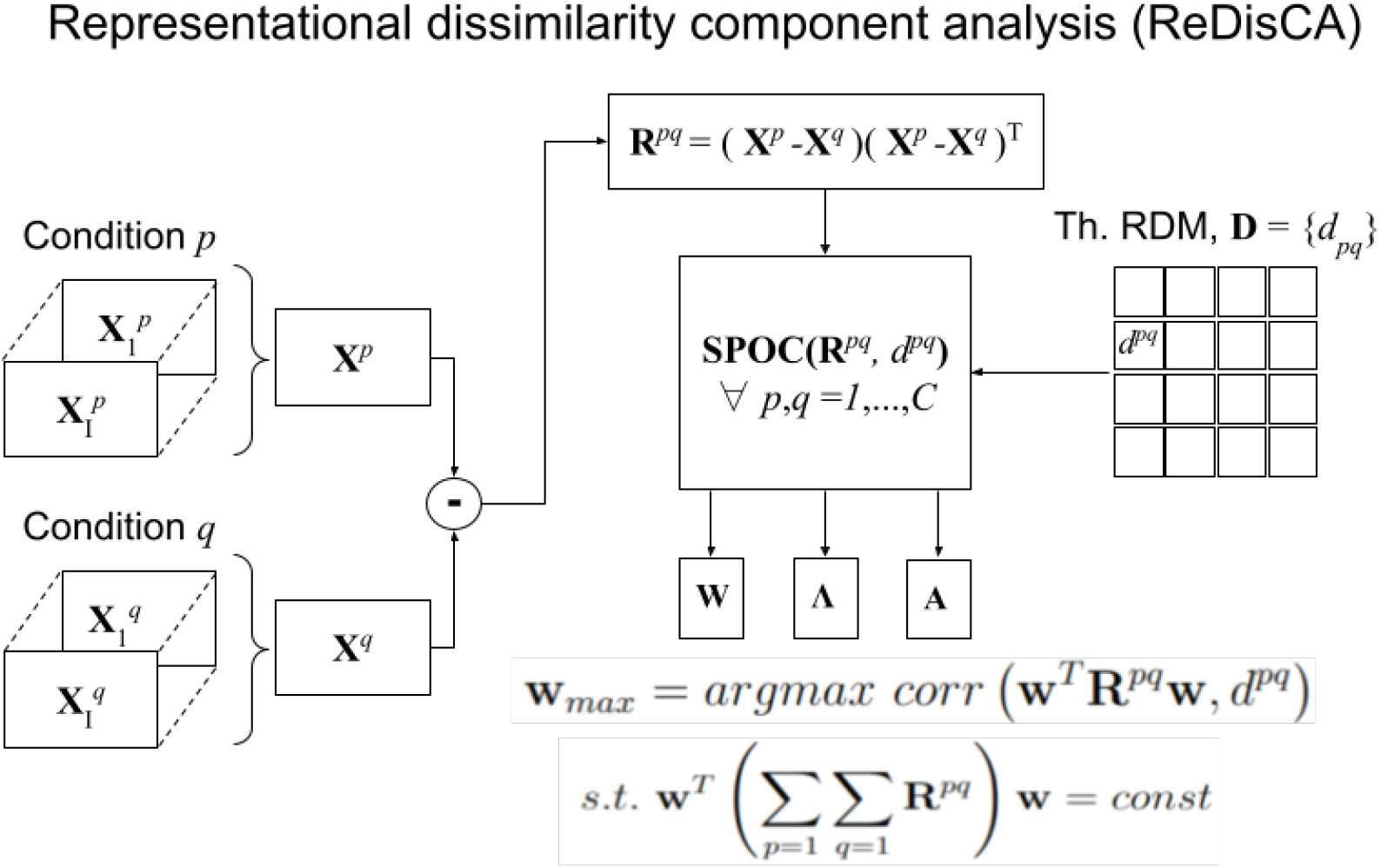
Representational dissimilarity component analysis (ReDisCA) diagram. The problem of finding the spatial components whose output ERP follow the desired RDM is reduced to SPoC covariance optimization. Instead of time-window covariance matrices **C**(*e*) and the associated values of behavioral variable *z*(*e*) observed for a set of latency values *e* = 1, … *E* ReDisCA uses activation difference covariance matrices **R**^*pq*^ and the elements *d*^*pq*^ of the theoretical RDM correspondingly, see also Table 1.

**Figure 3:**
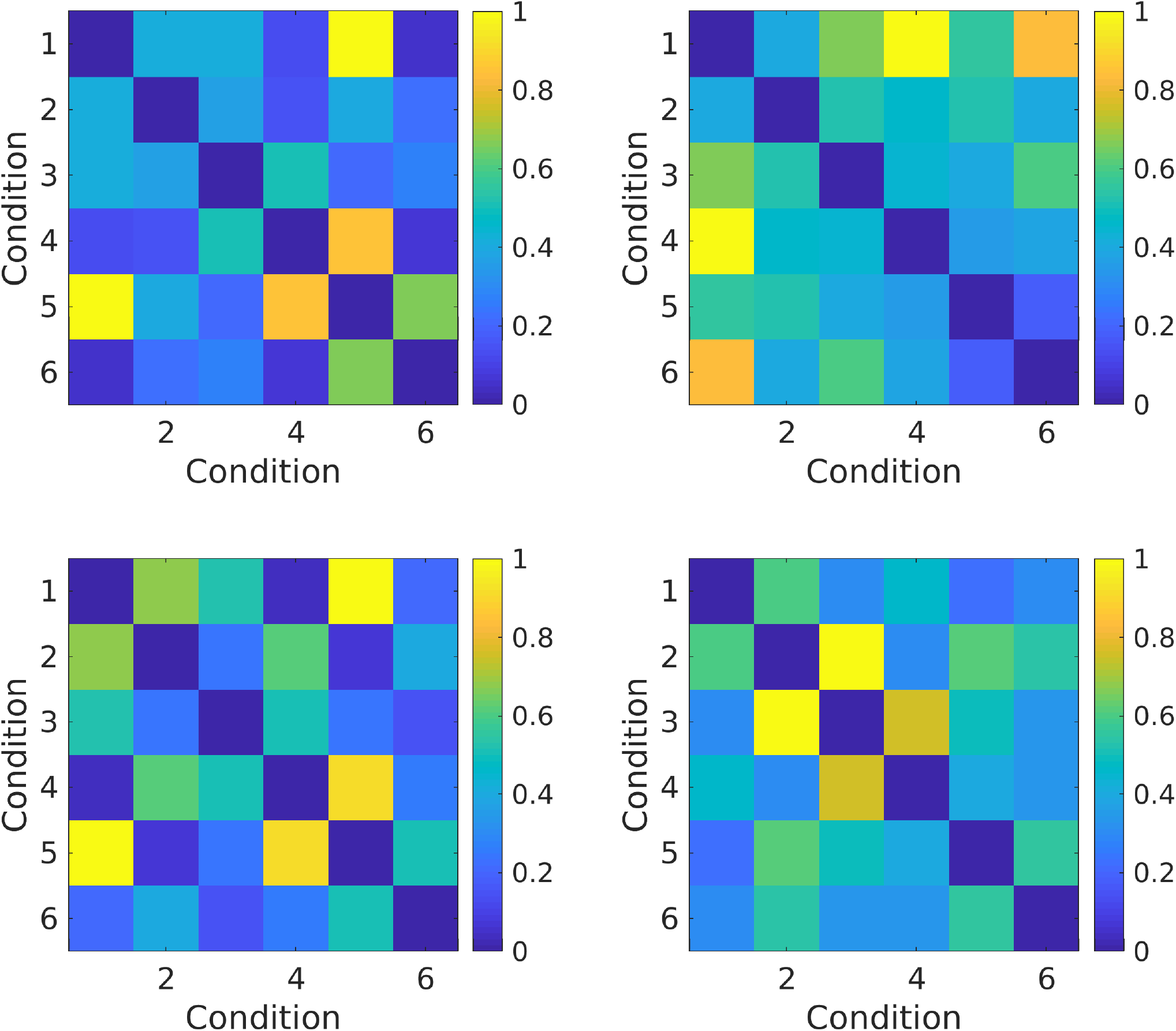
Theoretical RDM matrices for each of the four simulated sources generated within an exemplar Monte-Carlo trial.

### 3.1. ROC curves

Within the first set of simulations we evaluated the methods’ performance using area under ROC curve (ROC AUC) metrics. To compute the ROC curves we first determine the counts of true positives (TP), false positives (FP), true negatives (TN), and false negatives (FN).

To this end as described in Section 2.4 at each *k*-th Monte-Carlo iteration corresponding to a neuronal source placed at a randomly chosen location **r**_*k*_ we use the scans *ρ*_*m*_, *m* = 1, …, *M* described in Section 2 and for each threshold value *θ* we find a set of indices *ℳ* _*>*_ of cortical vertices *m* for which *ρ*_*m*_ ≥ *θ*. Then, we count the number of vertices in *ℳ* _*>*_ that fall within a sphere with *r*_*max*_ = 0.01 m centered around the true source location **r**_*k*_ simulated during the *k*-th Monte-Carlo iteration and save it to the corresponding true positives count array *TP*_*k*_(*θ*). The count of vertices from *ℳ* _*>*_ that fall outside the sphere with radius *r*_*max*_ is added to the false-positives count array *FP*_*k*_(*θ*).

Next we find a complementary set *ℳ* _*<*_ of cortical vertex indices *m* such that *ρ*_*m*_ *< θ*. We then count the number of vertices from *ℳ* _*<*_ that appear within the sphere with *r*_*max*_ = 0.01 m centered around the true source location **r**_*k*_ and save it to the corresponding false negatives count array *FN*_*k*_(*θ*). The count of vertices from *ℳ* _*<*_ outside the sphere with radius *r*_*max*_ is saved to the true-negatives count array *TN*_*k*_(*θ*).

Then, for each threshold value *θ* we compute the average true positive rate (*TPR*(*θ*)) or sensitivity and the average false positive rate *FPR*(*θ*) equal to 1 − *specificity* as

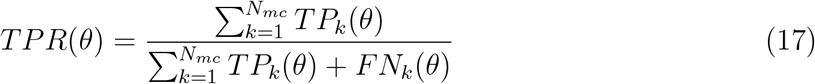

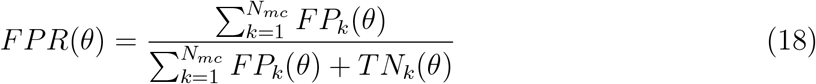

The pair (*FPR*(*θ*), *TPR*(*θ*)) parameterized by *θ* forms the ROC curve that we use to compare between different RSA approaches studied in this paper.

### 3.2. Performance gauges for realistic simulations

In the realistic simulations with multiple sources described in Section 2.4.3 we created *P* = 4 simultaneously active sources placed randomly on the cortex during each Monte-Carlo iteration, each with the specific theoretical RDM 𝔻^*p*^. In this case, we gauged the performance based on the distance between the simulated source locations and those that were estimated using ReDisCA and the two source space RSA implementations that performed best during the first set of simulations with a single source, see Section 2.4.2.

To compute this metric at each Monte-Carlo iteration we used the scans 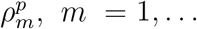, *M*, *p* = 1, …, *P* corresponding to the *p*−th source RDM 𝔻^*p*^ and identified the cortical vertex index *m*^*^ corresponding to the maximum of 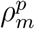 as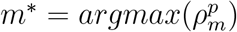. We then used the coordinates vector **r**^*^ of the *m*^*^ vertex and computed the distance between it and the true location of the *p*-th source 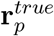, see Section 2.4.3 as 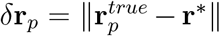. We then plotted the histograms *δ***r**_*p*_ observed in the Monte-Carlo trials for each of the compared methods.

At each Monte-Carlo iteration we have also computed the correlation coefficients between spatial topography vector 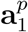 of the first ReDisCA component and the corresponding true source topography vectors 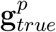, *p* = 1, …, *P*. Having performed a set of Monte-Carlo trials we summarized the observed correlation coefficient distributions in the form of histograms.

Finally, we have compared the target RDMs 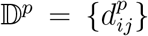 from the *p*-th source to the empirical RDMs 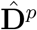. These empirical RDMs can be obtained from spatially filtered data using ReDisCA-derived filters **w**^*p*^ as 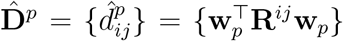, where **R**_*ij*_ is the *i*-th and *j* − *th* condition difference time series covariance matrix as described in section 2.3. Then, we measured the correlation coefficients between the vectors derived from the upper triangular elements of 𝔻^*p*^ and 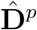 for each *p*−th source, *p* = 1, …, *P*. Using multiple Monte-Carlo trials, we summarized the distributions of these correlation coefficients in histograms.

## 4. Results

### 4.1. Simulations

In Figure 4, we illustrate the outcomes of the first set of our simulations where we compared ReDisCA with four versions of the source space RSA, see Figure 1 for their detailed diagram. As described before, the aim here was to detect a single neuronal source using a noisy version of its specific RDM. In Figure 4.a, the blue curve represents ReDisCA’s performance in the detection of the representational dissimilarity subspace using the averaged evoked response data. This method significantly outperforms four different versions of the source space RSA that employ two different inverse operators: the minimum norm estimator and the LCMV beamformer. Out of the four source space RSA approaches LCMV beamformer based on single trials (BF S.T.) approach appears to perform best. Importantly, ReDisCA yields better ROC AUC values in detecting spatial component linked to the targeted neuronal source without relying on electromagnetic inverse modeling and with significantly fewer computations as compared to the source space RSA. To make this conclusion and enable the comparison we employed the electromagnetic inverse modeling to align ReDisCA’s results with those of conventional RSAs that operate explicitly in the space of neuronal sources, see expressions (13) and 14). Figure 4.b illustrates noisy (blue) and noise-free (red) averaged ERP time series and the theoretical dissimilarity matrix used in simulations is shown at the bottom. Figures 4.c and 4.d show similar results but for SNR = 0.1. As we can see, in this atypically low SNR case ReDisCA’s performance only slightly deteriorates and remains superior to that of the other methods. ReDisCA appears capable of accurately locating the target source in nearly 85% of cases at almost zero false alarms rate. Comparing the curves of classical RSA approaches (MNE AV RSA, MNE S.T. RSA, BF AV RSA, BF S.T. RSA) across two SNR values (panels a) and c) in Figure 4), an interesting finding emerges. We notice an improvement in the performance of the source space RSA methods under low SNR conditions compared to higher SNR scenarios in this single-source simulation. Our analysis reveals that this seeming paradox arises due to a decrease in false positives in the traditional RSA outputs as SNR decreases. Despite the use of a realistic brain noise model with spatial correlations, the resulting data matrix appears ‘empty’. This results in the situation where despite the inverse modeling the sole persistent component of the single source remains spread across numerous cortical vertices in the source time series estimates, which leads to a surplus of high correlations between the observed and theoretical RDMs. The increased noise levels, however, rectify this issue, rendering RDM correlation scans more specific which in turn improves the corresponding ROC curves.

**Figure 4:**
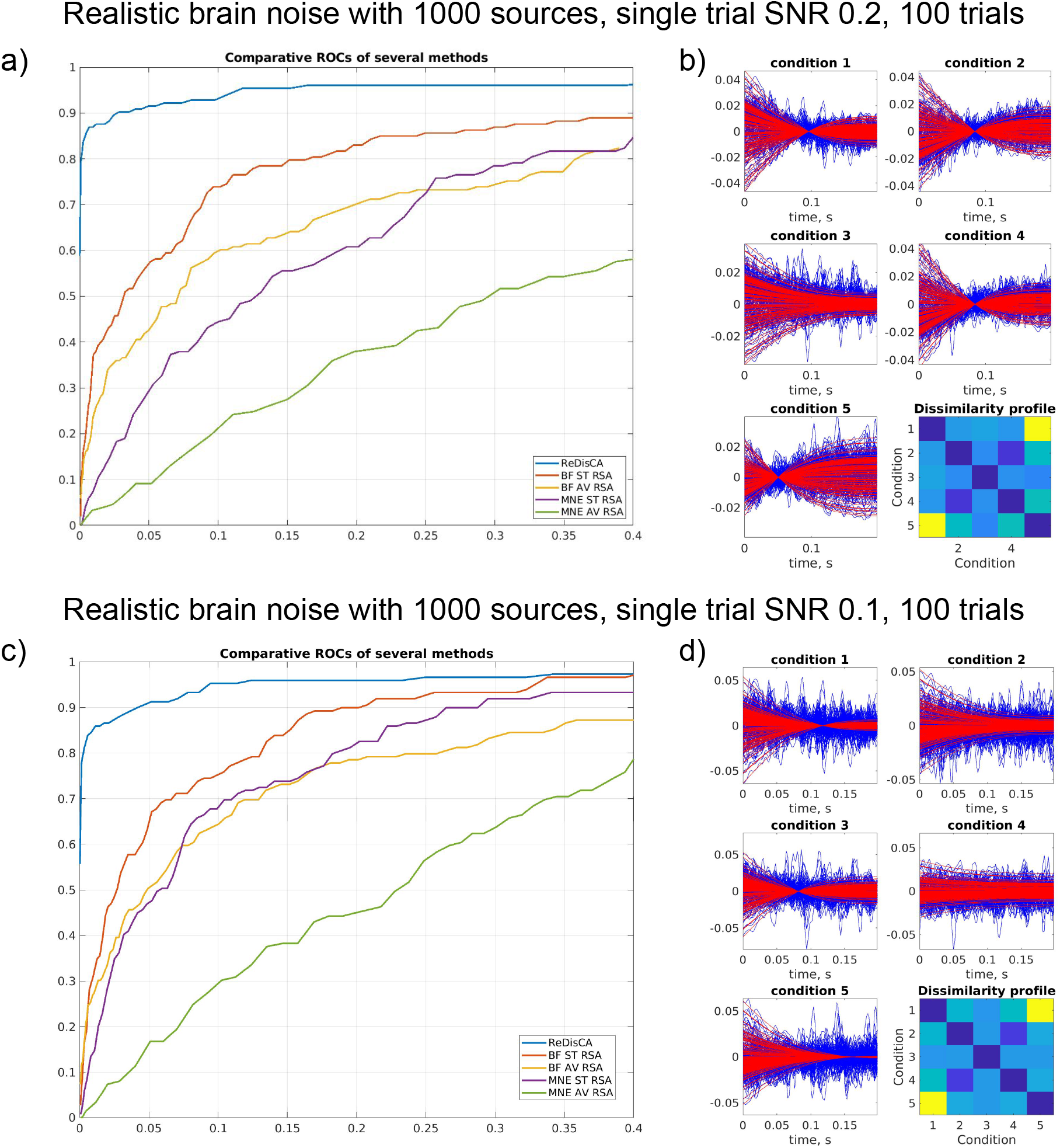
Results of 100 MC trials of simulations in single source detection scenario. a) and c) the ROC curves for ReDisCA and four source space RSA versions, see Section 2 for two different SNR levels. b) and d) - noisy (blue) and noise-free (red) averaged ERP time series for the two SNR values and the theoretical dissimilarity matrix used in the simulations.

This observation aligns with the lower ROC-AUC values observed in the source space RSAs based on the averaged ERP data (BF AV and MNE AV) compared to their single-trial counterparts (BF S.T. and MNE S.T.). In the latter case, dissimilarity scores are computed on individual trials before averaging, while in the former the averaging precedes computation of the vertex-specific between-condition dissimilarity scores based on the inverse modeling of these averaged ERP data (refer to Figure 1). This paradoxical behavior, however, does not manifest in more realistic simulations involving multiple cortical sources, as described next.

Next we describe the results of more realistic simulations where at each Monte Carlo trial we randomly seeded *M* = 4 sources each with its own RDM, see Section 2.4.3 for details. The results from 100 Monte Carlo trials of these realistic simulations for *C* = 5 conditions are presented in Figure 5 for two SNR values.

**Figure 5:**
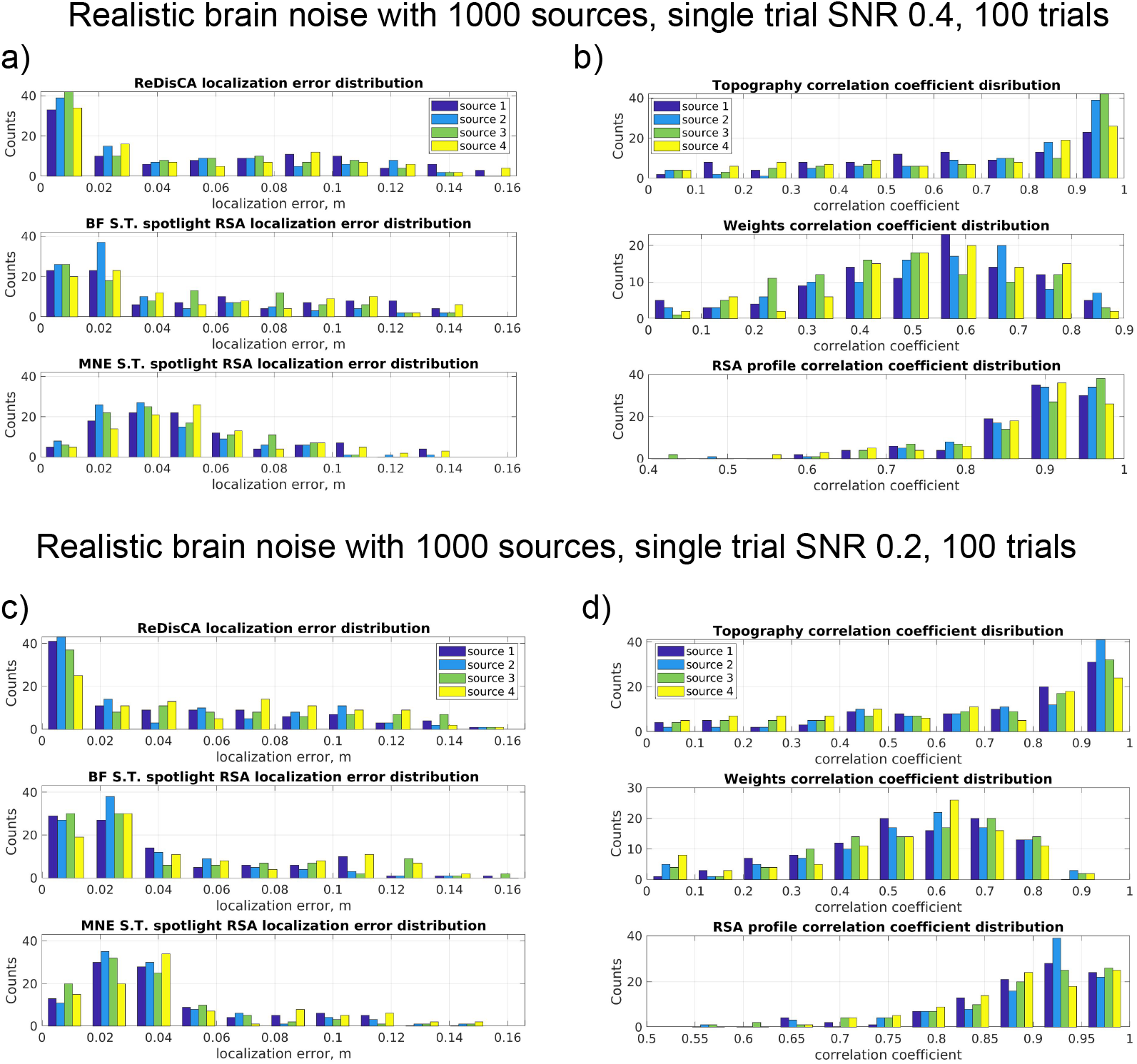
Results of 100 MC trials of realistic simulations with *M* = 4 randomly seeded sources and *C* = 6 conditions: a) Distribution of the true topography and ReDisCA derived pattern correlation coefficient, b) Distribution of the true topography and ReDisCA spatial filter weight vector correlation coefficient, c) Distribution of the correlation coefficient of true RDM and RDMs derived from the multichannel data with the first ReDisCA derived spatial filter, d) Distribution of ReDisCA source localization error, e) Distribution of the MNE spotlight RSA source localization error.

In Figure 5.a we show the distribution of source localization errors for four sources (different colors) for the three methods: ReDisCA, BF S.T. RSA, and MNE S.T. RSA. Out of compactness considerations we have chosen to present here the results of only two best-performing source space RSA methods, BF S.T. and MNE S.T. As one can see ReDisCA’s distribution appears to have the largest proportion of cases where all four sources are localized with errors below 1 cm. The distributions of source localization errors delivered by BF S.T. and MNE S.T. are increasingly shifted to the right and result in greater median source localization error as compared to ReDisCA.

Panel b) shows ReDisCA-specific metrics described in Section 3.2. From top to bottom we show the distributions of 1) topography correlation coefficient between ReDisCA identified **a**_1_ and the true simulated source topography 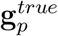, 2) ReDisCA weights **w**_1_ and the true topography 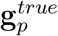 correlation coefficient and finally 3) correlation coefficient between the target 𝔻^*p*^ and observed RDMs 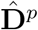 for *p* = 1, …, 4.

Since the spatial filters derived by ReDisCA get not only tuned to the target source but also attempt to tune away from the interfering sources [17, 48] the true source topographies appear to be much better aligned with ReDisCA derived topographies (patterns) than with the corresponding weight vectors. As evident from the bottom plot in Figure 5.b the observed RDMs appear to be well correlated with the target RDMs for all four sources. Panels c) and d) of Figure 5 show similar plots but for the decreased Signal-to-Noise Ratio (SNR) equal to 0.2, allowing us to observe that ReDisCA remains operable in these harsh conditions. It is important to note that in simulations by combining noiseless sensor data with noise data matrices, see equation (16), we effectively control the SNR based on the ratio of root mean powers between these two matrices. Therefore, as the number of sources within the noiseless data matrix increase, the SNR per individual source automatically decreases. Hence, in the case of four sources, the SNR of 0.2 represents a notably more challenging task compared to a single-source scenario with the SNR of 0.1.

Finally, we explore the dependence of localization error on the number of conditions employed in RSA analysis. It is intuitive to expect that the increase in the number of conditions should lead to improved source localization performance in the classical RSA scenario and better identification of the representationally relevant components by ReDisCA. As before, to align ReDisCA’s results with those of source space RSA approaches we fitted a dipole to ReDisCA-derived source topography vectors 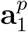, *p* = 1, …, 4. The results are presented in Figure 6 in the form of three graphs corresponding to the average median localization error achieved by the three techniques. As expected the increase in the number of conditions leads to the decrease of source localization error. ReDisCA furnishes the best performance for all *C*, the mean median error appears to be less than 2 cm for *C* = 6 conditions.

**Figure 6:**
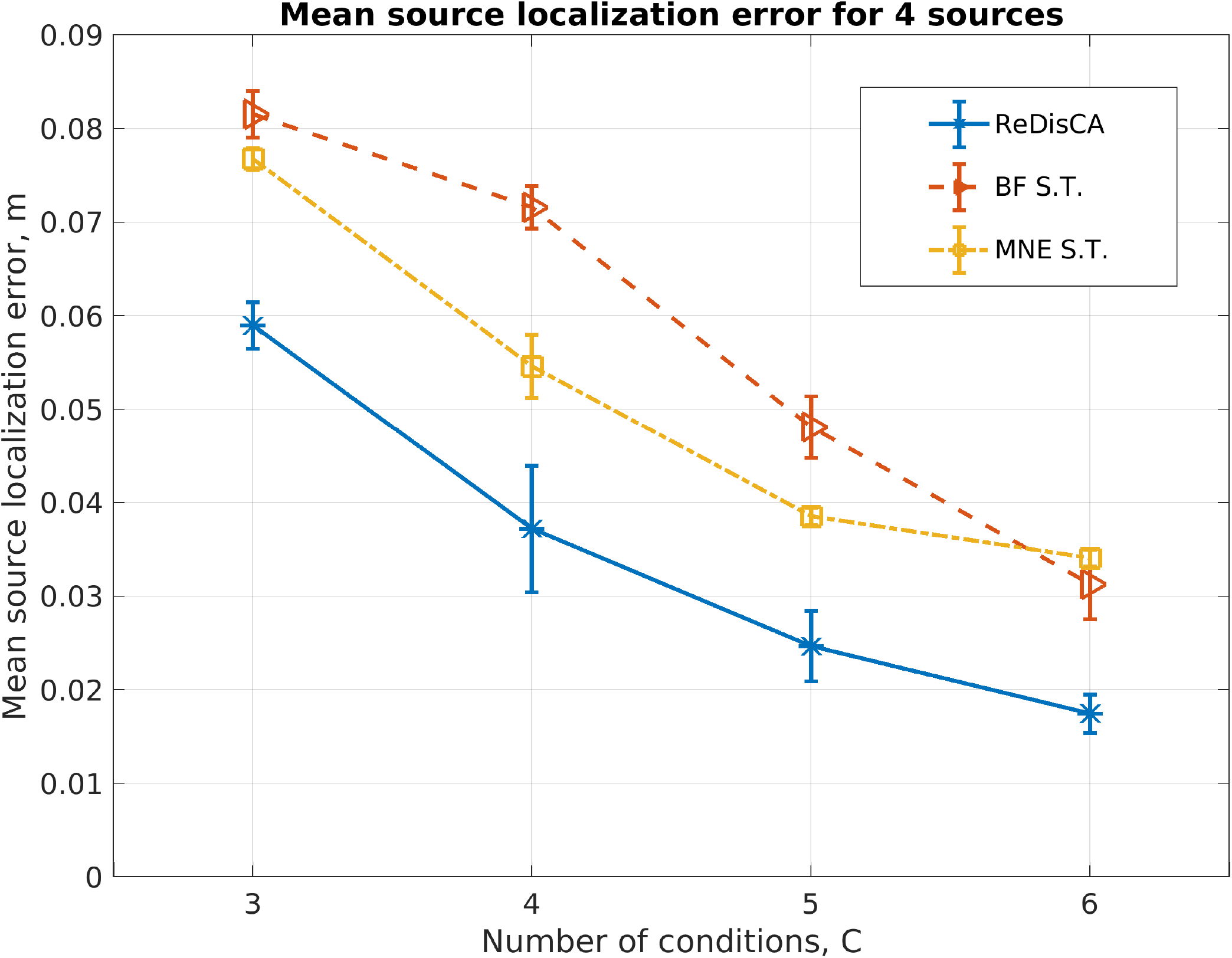
Average median localization error of 4 simultaneously active sources (each with its own RDM) as a function of the conditions count *C* for three methods ReDisCA, beamformer-powered single trial RSA, and MNE-powered single trial RSA. As expected the increase in the number of conditions leads to the decrease of source localization error. ReDisCA furnishes the best performance for all *C*, the mean median error appears to be less than 2 cm for *C* = 6 conditions.

### 4.2. Real data analysis examples

In this section we demonstrated the use of ReDisCA for analysis of two real-life datasets. The first dataset is a publicly available low-density EEG N170 dataset available at https://osf.io/pfde9/ and described in [22]. In that data the N170 was elicited in a face perception task with stimuli from [51]. In this task, an image of a face, car, scrambled face, or scrambled car was presented on each trial in the center of the screen, and participants responded whether the stimulus was an “object” (face or car) or a “texture” (scrambled face or scrambled car). The data were preprocessed and three ICA components corresponding to ocular and cardiac artifacts were removed from the data. Then, the ERP were computed by averaging responses within each of the stimulus types. The dataset comprises the EEG data from 40 subjects. In our analysis for demonstration purposes we used single subject data recorded from the first participant with index “1”.

The second dataset is an Elekta Neuromag 306 MEG dataset from the first run of the RSA study by [27]. In this open-access article, the authors employed source space RSA and demonstrated distinct differences in the timing, brain regions involved, and dynamics of visual processing of faces and tools during the categorization stage. They found that face-specific spatiotemporal patterns were linked to bilateral activation of ventral occipito-temporal areas during the feature binding stage at 140–170 ms. In contrast, tool-specific binding-related activity was observed within the 210–220 ms window, located in the intraparietal sulcus of the left hemisphere. Brain activity common to both categories began at 250 ms and included widely distributed assemblies within the parietal, temporal, and prefrontal regions. A more detailed description of the spatial-temporal dynamics can be found in Figure 3 of [27]. In our analysis we used the data recorded from the first subject labeled as “AD”.

#### 4.2.1. ReDisCA of a low-density EEG dataset

We first applied ReDisCA to explore the spatial-temporal structure of the response to meaningful versus meaningless stimuli that were formed as the scrambled versions of the original images. For that, we formed the theoretical RDM shown in Figure 7.a. Then, scanning over time windows of duration *T* = 150 ms and calculating the *p*-values we obtain their color-coded map shown in Figure 7.b with rows corresponding to the components and columns encode time. We then visualized uncorrected *p*-values corresponding to the first component and identified the time interval corresponding to a continuous segment of *p <*0.05. As can be seen from Figure 7.d we found such a segment at around *t* = 400 ms. The corresponding pattern is visualized in panel c) and has highly pronounced occipital topography. This result is in agreement with observations made in a high-density EEG-based study [37] that demonstrates a statistically significant difference between meaningful and meaningless stimuli occurring in the time window around 300-500 ms and is localized to the occipital area of the scalp. ReDisCA missed the first peak at around 100 ms reported by [37] but that could be explained by a smaller number of electrodes in the current dataset as compared to the one analyzed in [37]. At the same time, our analysis of an MEG dataset, see Figure 15 in Section 4.2.2, shows a significantly different activation of ReDisCA component starting at 160 ms and contrasting the meaningful vs. meaningless visual stimuli. This component has dominantly occipital topography.

**Figure 7:**
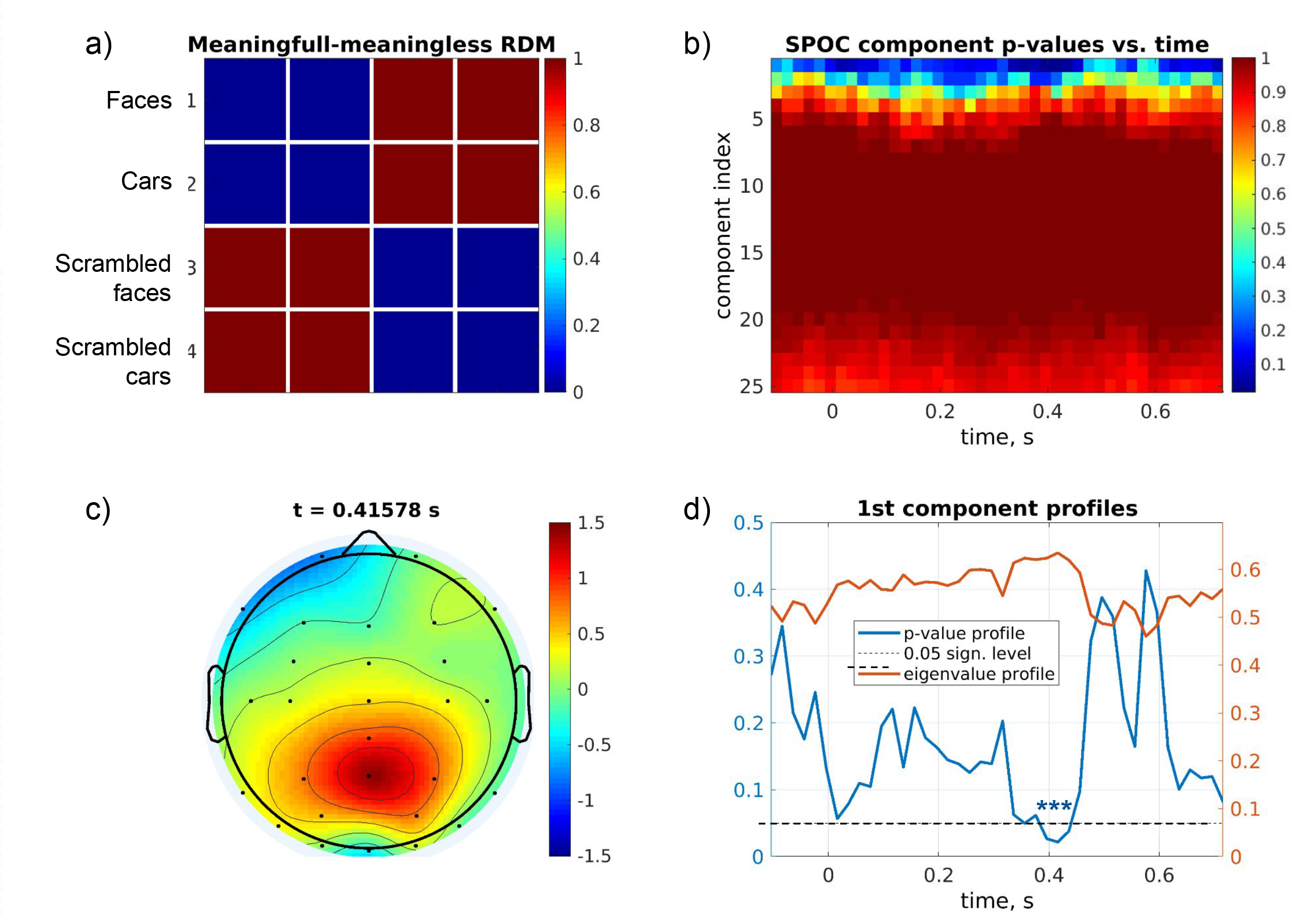
Meaningful vs meaningless stimuli perception analysis. a) Theoretical RDM contrasting the two types of stimuli. b) Color-coded map of component *p*-values, each row corresponds to a component and columns encode time. c) Topography (pattern) of the first and the only significant component during the time interval around *t* = 400 ms. d) Temporal profiles of the uncorrected *p*-values corresponding to the first component. The significant segment is marked with asterisks.

A significant component aligned with the theoretical meaningful vs. meaningless RDM was observed over 3 adjacent time windows. Figure 8 shows the discovered topographic patterns of the first ReDisCA component, the corresponding time courses of this component observed within the four different conditions as well as the observed (or empirical) color-coded RDM in the bottommost panel. As we can see the yellow and violet timecourses corresponding to the meaningless stimuli appear disentangled from the red and blue ones observed during the presentation of meaningful pictures. The topography of the first significant ReDisCA component (*p <* 0.05) remains highly similar over the three consecutive time windows. Interestingly, the time series of this ERP component observed during the presentation of faces (blue) exhibits longer-lasting traces as compared to that corresponding to the activity during the presentation of cars (red curve) which is consistent with the findings reported in several studies exploring neural correlates of face perception in humans [18]. Consequently, the observed RDMs indeed appear to closely resemble the target RDM, see Figure 7.a used for the described inquiry. Comparing the evolution of the topography of the first (the only one found significant) ReDisCA component we can observe its gradual displacement in the sagittal plane moving downwards with time followed by deepening (the topography widens) of the source at the last time stamp. This may correspond to the traveling wave patterns found in the visual area [52, 4].

**Figure 8:**
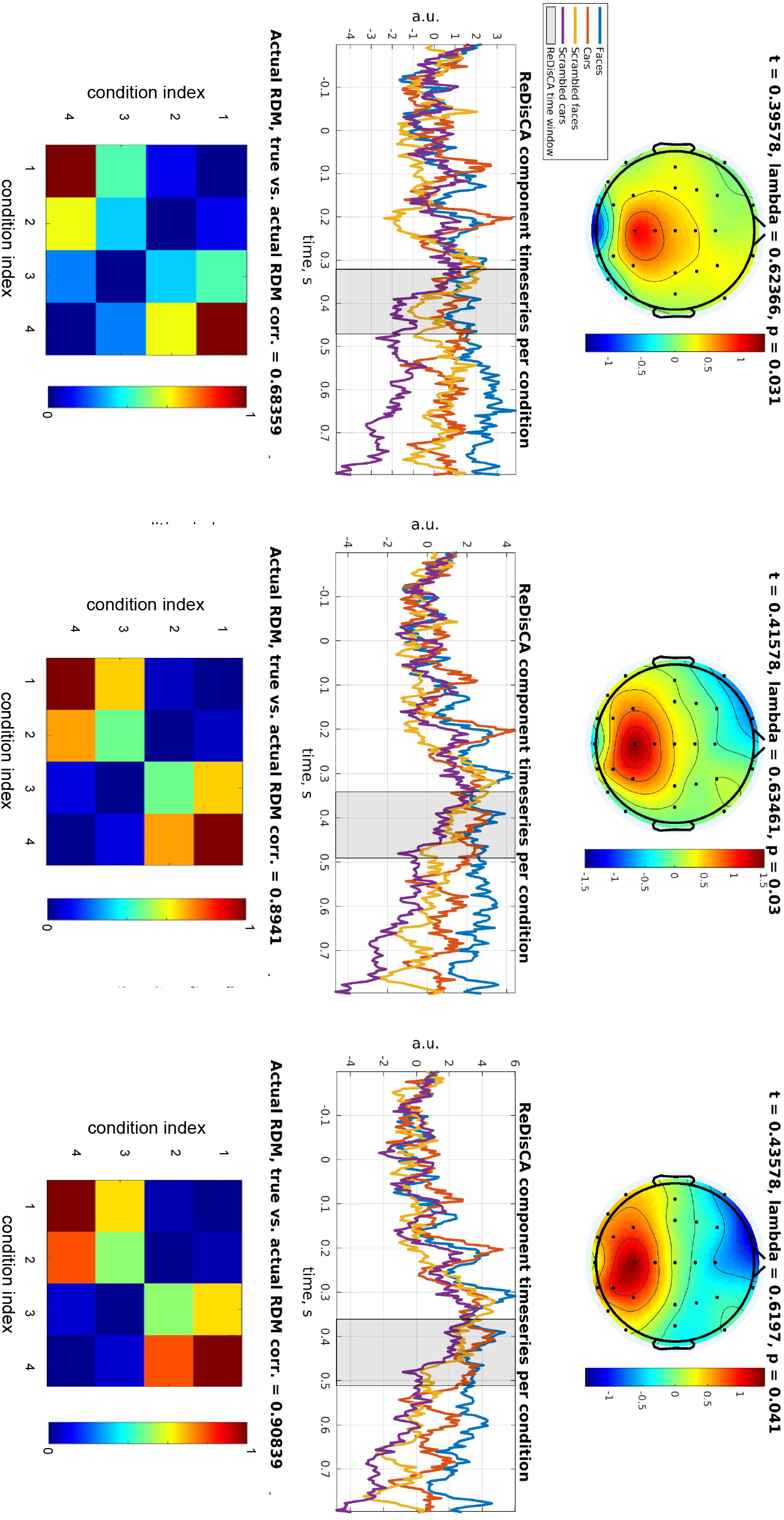
Meaningful vs. meaningless stimuli perception analysis. Topography, spatially filtered timecourses and the observed RDMs for the three consecutive time slices at around *t* = 400 ms, see also Figure 7.d

We proceeded by examining the activity associated with particular meaningful stimuli using the RDMs displayed in Figure 9.a for faces and Figure 9.b for cars, respectively. These RDMs guide our search towards a component with a distinct activity in one condition (face or car) different from that observed in the remaining three conditions. Additionally, unlike what may be the case in the decoding-based multivariate pattern analysis approaches, the RDMs prescribe the activity during the remaining three conditions to be similar. Although a classifier can be built to enforce a compact representation within each of the classes, the RSA offers a greater flexibility in imposing the geometric constraints on the discovered activity using non-binary RDM matrices. See the analysis reported in Figures 16 and 17 in Section 4.2.2.

**Figure 9:**
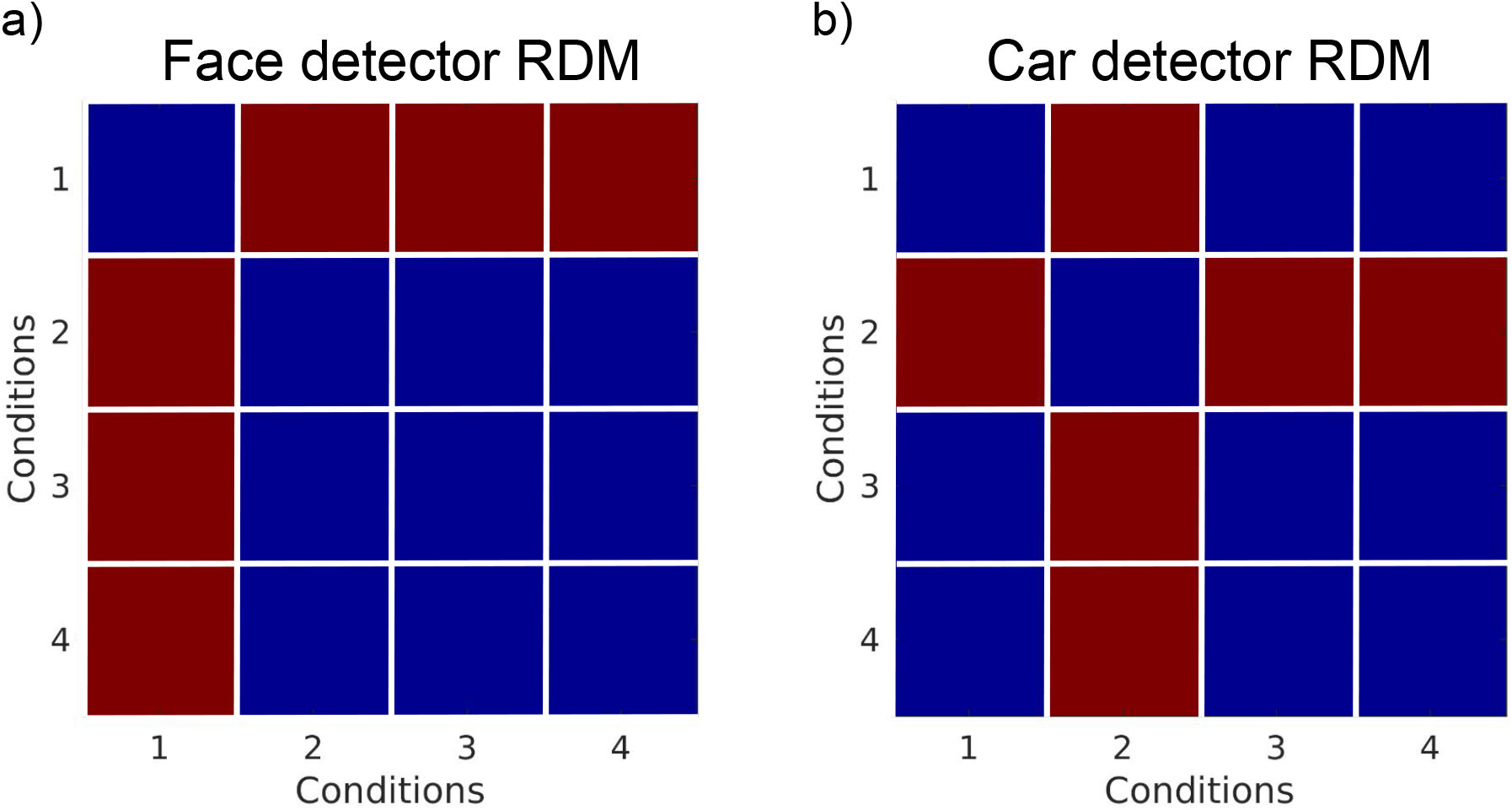
Theoretical RDMs corresponding to faces (a) and cars (b)

According to the large volume of studies N170 ERP component is considered to be face specific and occurs at around 170 ms latency. We have therefore selected the time window of duration *T* = 100 ms centered at 200 ms and applied ReDisCA to it using the theoretical RDM shown in Figure 9.a. Figure 10 shows ReDisCA’s output and as we can see from the top panel of 10 the first and the only significant ReDisCA component has topography corresponding to a source in the right fusiform gyrus, the area known to be crucial to perception of facial information and differentially activated in response to regular or scrambled face stimuli [16, 27].

**Figure 10:**
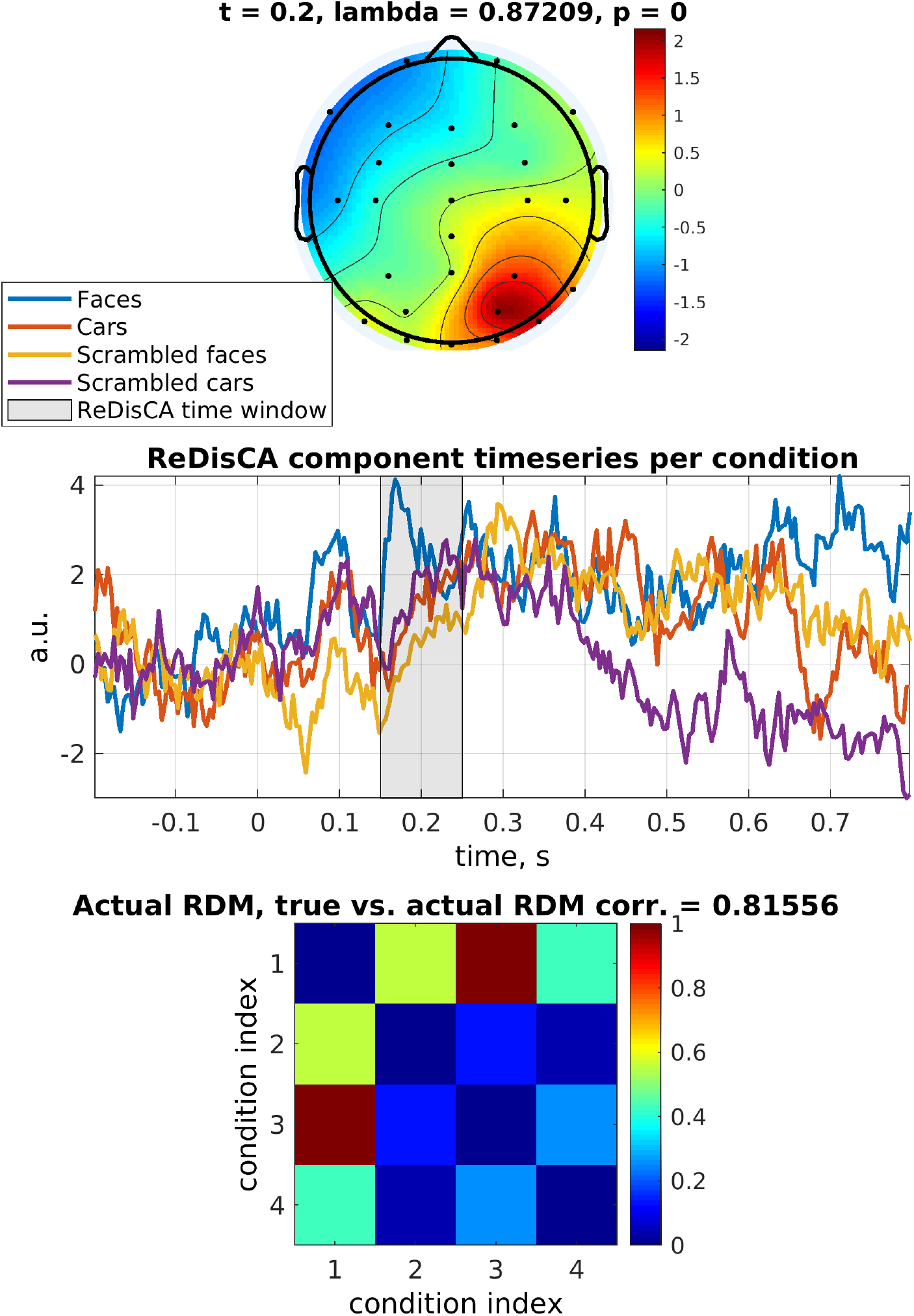
ReDisCA of face-specific responses using the RDM depicted in Figure 9.a. The top panel shows the topography (pattern) of the first and the only significant ReDisCA component. The topography corresponds to a source in the right fusiform gyrus, the area pivotal to the perception of facial information. In the central panel the corresponding ERP components in the four conditions are displayed. The observed RDM is shown in the bottommost panel and exhibits a high correlation coefficient of 0.82 with the theoretical RDM.

The corresponding ERP components obtained by filtering the multichannel ERP with a spatial filter derived by ReDisCA are presented in the panel below the topography. We indeed can observe a burst of activity around 170 ms in the “face” condition while in the other three conditions the response curves do not have such a burst and are very closely aligned implementing the requirement imposed by the RDM matrix. In agreement with other studies, the discovered face-related ERP components remain active over a long duration of response. The observed RDM is shown in the bottommost panel. We can see that it aligns well with the theoretical RDM and exhibits a high correlation coefficient of 0.82.

Finally, we performed similar processing but using the theoretical RDM from Figure 9.b designed to highlight response components specific to the processing of car images. We have found two significant components with *p <* 0.01 shown in Figure 11. The first components appear to have a lower occipital topography with the time course exhibiting deflection at around 150 ms that is specific to the second condition (car images). This reflects activity in the ventral visual pathway during the image perception task. The traces of this ReDisCA component appear to be remarkably similar in the other three conditions not only during the target interval but also over the entire response duration. The observed and the theoretical RDM appear highly correlated with a correlation coefficient greater than 0.99.

**Figure 11:**
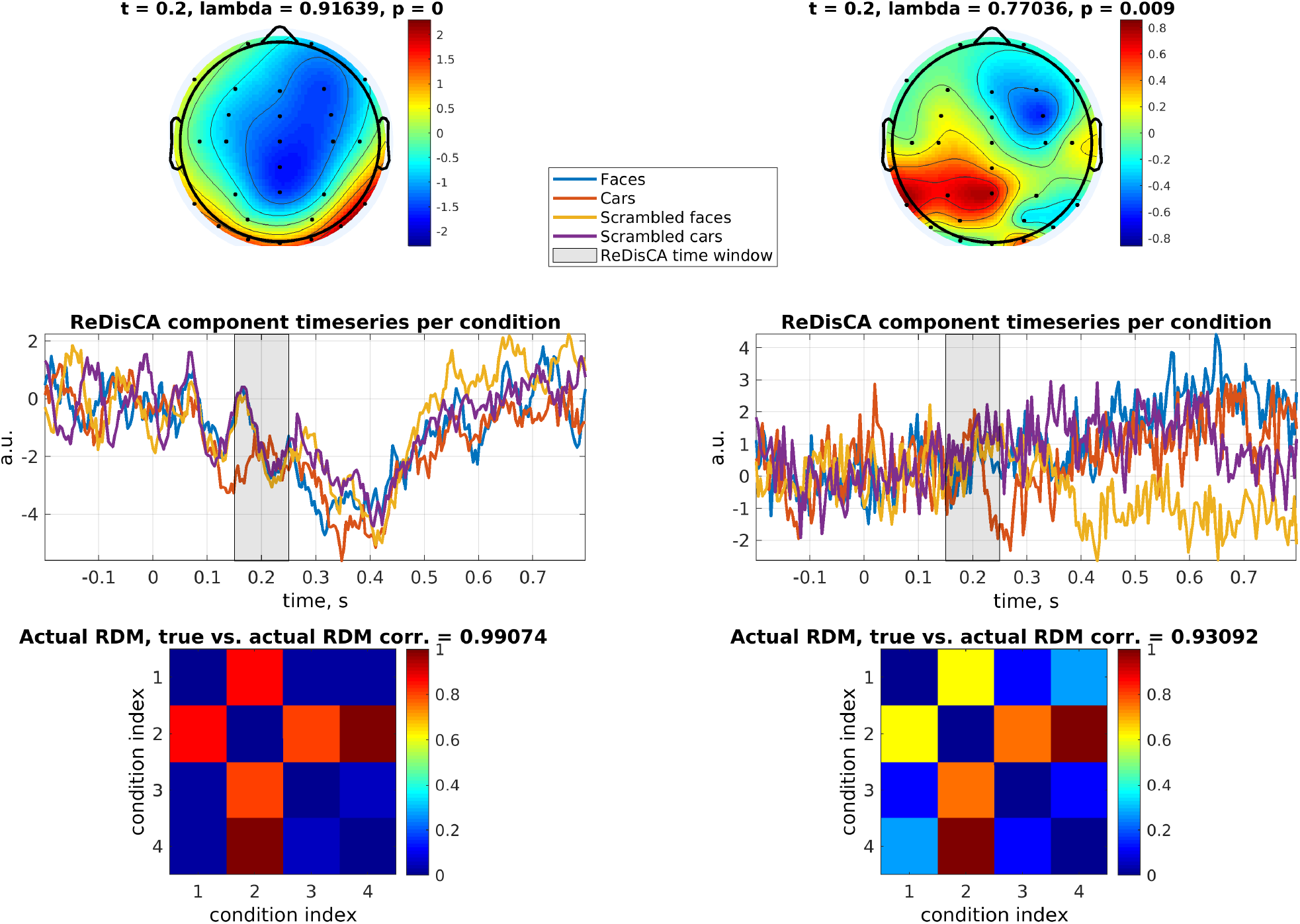
Two significant components discovered by ReDisCA using car detector RDM (Figure 9.b) applied at *t* = 170 ms latency.

Interestingly, the second component has a pronounced topography that may reflect the activity in the dorsal visual pathway that is known to accompany visual perception of objects and is hypothesized to reflect the neural processes of formation of spatial relations between the objects within the scene that is then fed to the ventral visual stream [3]. It may also be related to identifying the car as a graspable tool as far as the steering wheel is concerned [2].

Based on the above we can conclude that the reported real data analysis results obtained with a direct application of ReDisCA to the averaged ERP data appear to align well with the existing knowledge in the field of neuroimaging related to visual perception of faces and cars and differentiation between meaningful and meaningless stimuli including not only spatial but also temporal structure of this process.

These findings obtained with ReDisCA appear to be in line with what’s already known in the field of neuroimaging regarding the brain processes underlying visual perception. This includes distinguishing between meaningful and meaningless stimuli, delineation of face-specific and car-specific activation considering both the spatial and temporal aspects of this cognitive process.

#### 4.2.2. ReDisCA of the MEG visual stimuli categorization dataset

To enable a comparison to a more traditional source space RSA we applied ReDisCA to the MEG dataset from [27] corresponding to the first run of their experiment.

Similarly to [27], we divided the responses within each category into two equal parts and labeled them numerically as 1 and 2, resulting in total 6 subcategories labeled “face 1”, “face 2”, “tool 1”, “tool 2”, “nons. 1”, and “nons. 2”. In each of the subcategory we had 80 epochs and 480 epochs in total. Since ReDisCA operates on the averaged evoked responses, we averaged the single-trial responses to obtain six evoked response field (ERF) matrices, one for each subcategory. Each ERF response is a 204 *×*1500 matrix, reflecting the activity of 204 planar gradiometers over 1500 ms, with 500 ms of pre-stimulus and 1000 ms of post-stimulus intervals.

We have then applied ReDisCA to these six ERFs aiming to elucidate the components supporting cortical processing of each of the 3 categories. We used the theoretical RDM matrices shown in Figure 12 for each of the three categories. This time we applied ReDisCA to the entire 1500 ms time-window at once. We then used the spatial filters derived by ReDisCA and computed the time series associated with each of the representation dissimilarity components. In our presentation here we considered only the first three statistically significant ReDisCA components.

**Figure 12:**
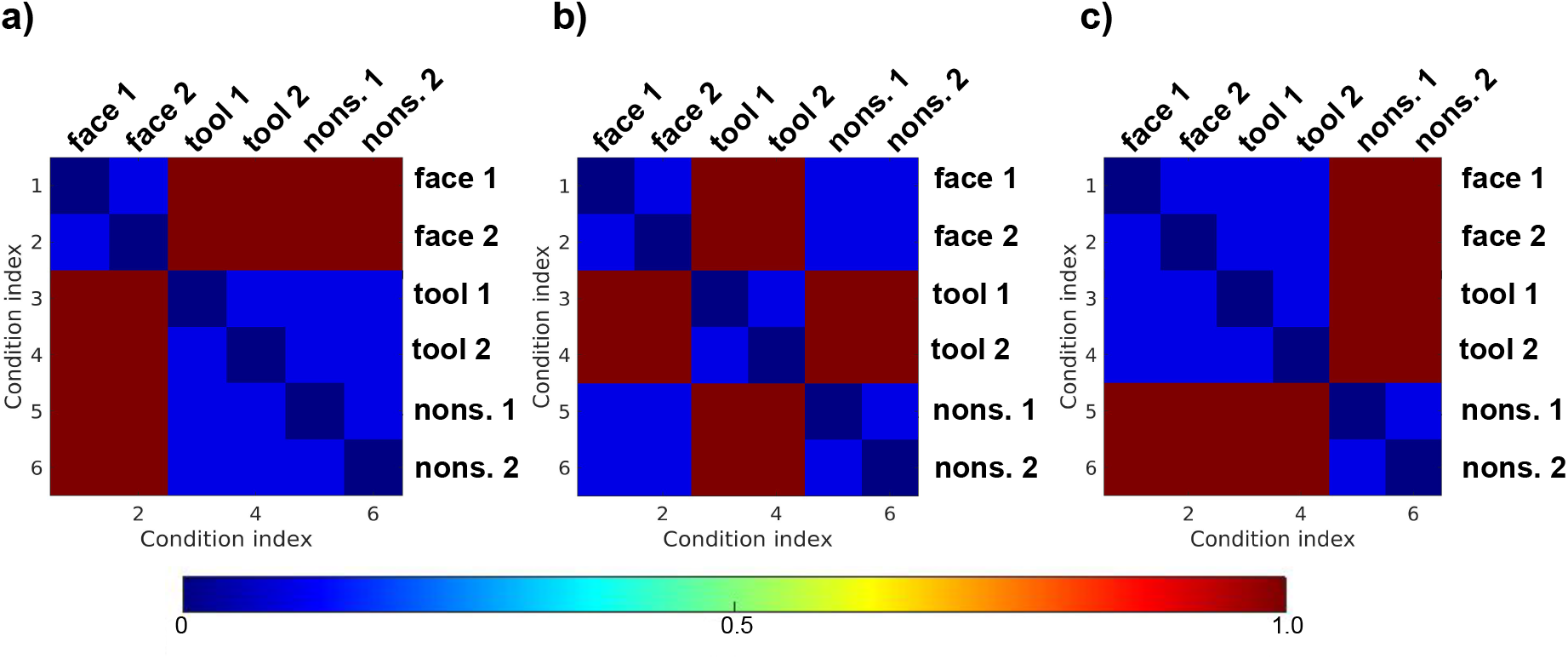
Three theoretical RDM matrices used for analysis of [27] dataset.

The top row of Figure 13 shows the temporal profiles for each of the 6 categories (“face 1”, “face 2”, …, “nons. 2”, see the legend) for the face-specific theoretical RDM from Figure 12.a. Panels a)-c) of Figure 13 correspond to the first three ReDisCA components. The title of each panel contains the component’s *p−* value. The bottom row of plots in Figure 13 depicts spatial patterns associated with each ReDisCA component discovered using the face-specific theoretical RDM.

**Figure 13:**
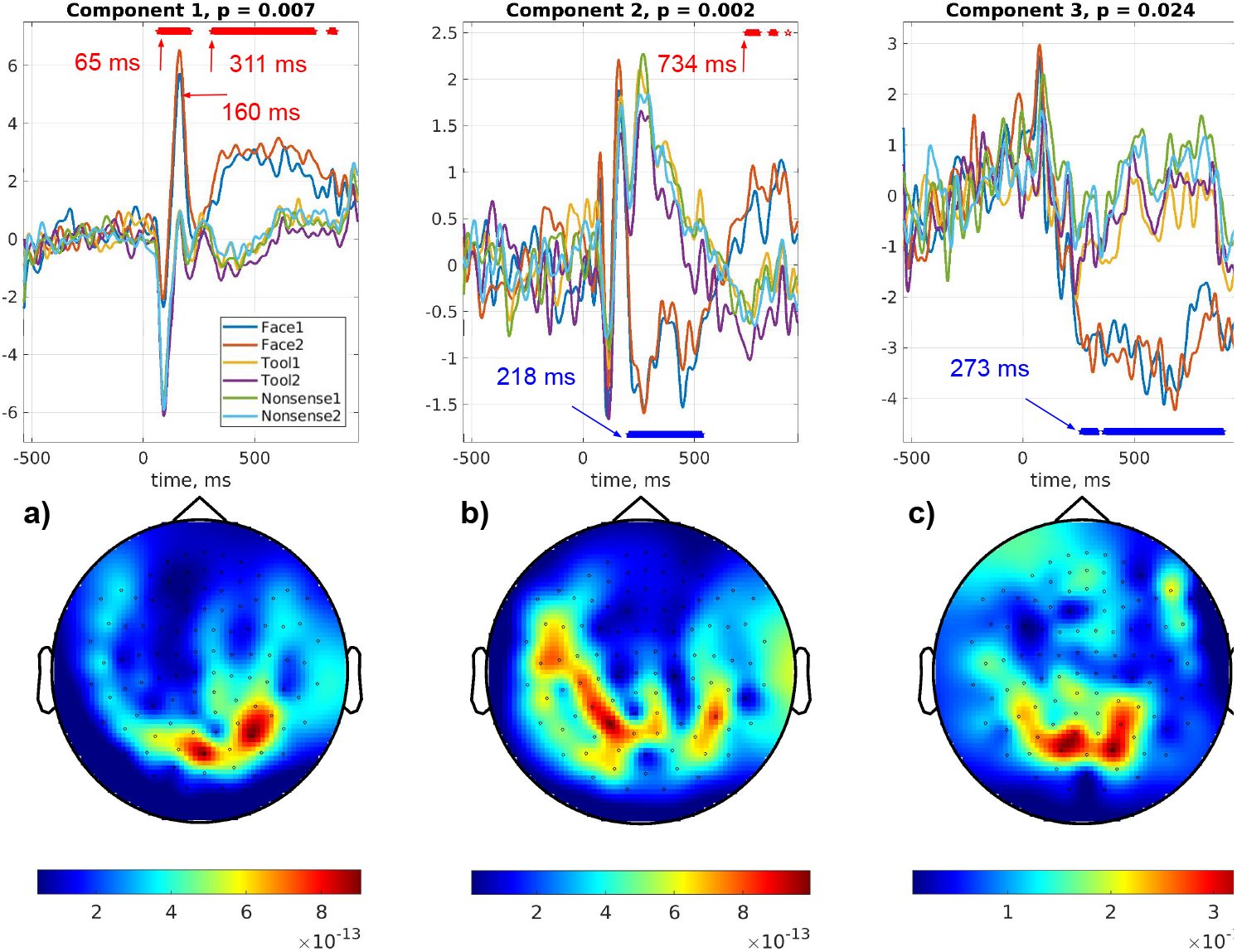
ReDisCA applied to the data from [27] with the face-specific theoretical RDM, see Figure 12.a. Panels a)-c) correspond to the first three ReDisCA components. Within each panel the top plot shows temporal profiles for each of the 6 subcategories (“face 1”, “face 2”, “tools 1”, “tools 2”, “nons. 1” and “nons. 2”), see the legend and the title for the associated *p*-value. The bottom plot shows the component’s spatial pattern.

To determine the intervals of significant difference in component activation time series between categories, we performed randomization tests. This involved permuting the sub-category labels of individual epochs, computing surrogate averages, and applying the corresponding spatial filters. We then corrected for multiple comparisons using the family-wise error rate (FWER) principle operationalized by the maximum statistics computed over the entire time interval. The results of the statistical testing are shown with red and blue asterisk lines located above and below the time series plots in the top panels of Figures 13 - 15. For the reader’s convenience and to facilitate a comparison of ReDisCA results with the source space RSA findings reported in [27], we have also indicated the starting times of the major significance intervals in each of the time series plots.

Similarly we have applied ReDisCA using the other two theoretical RDMs from Figures 12.b and 12.c and generated Figures 14 and 15 with ReDisCA results.

**Figure 14:**
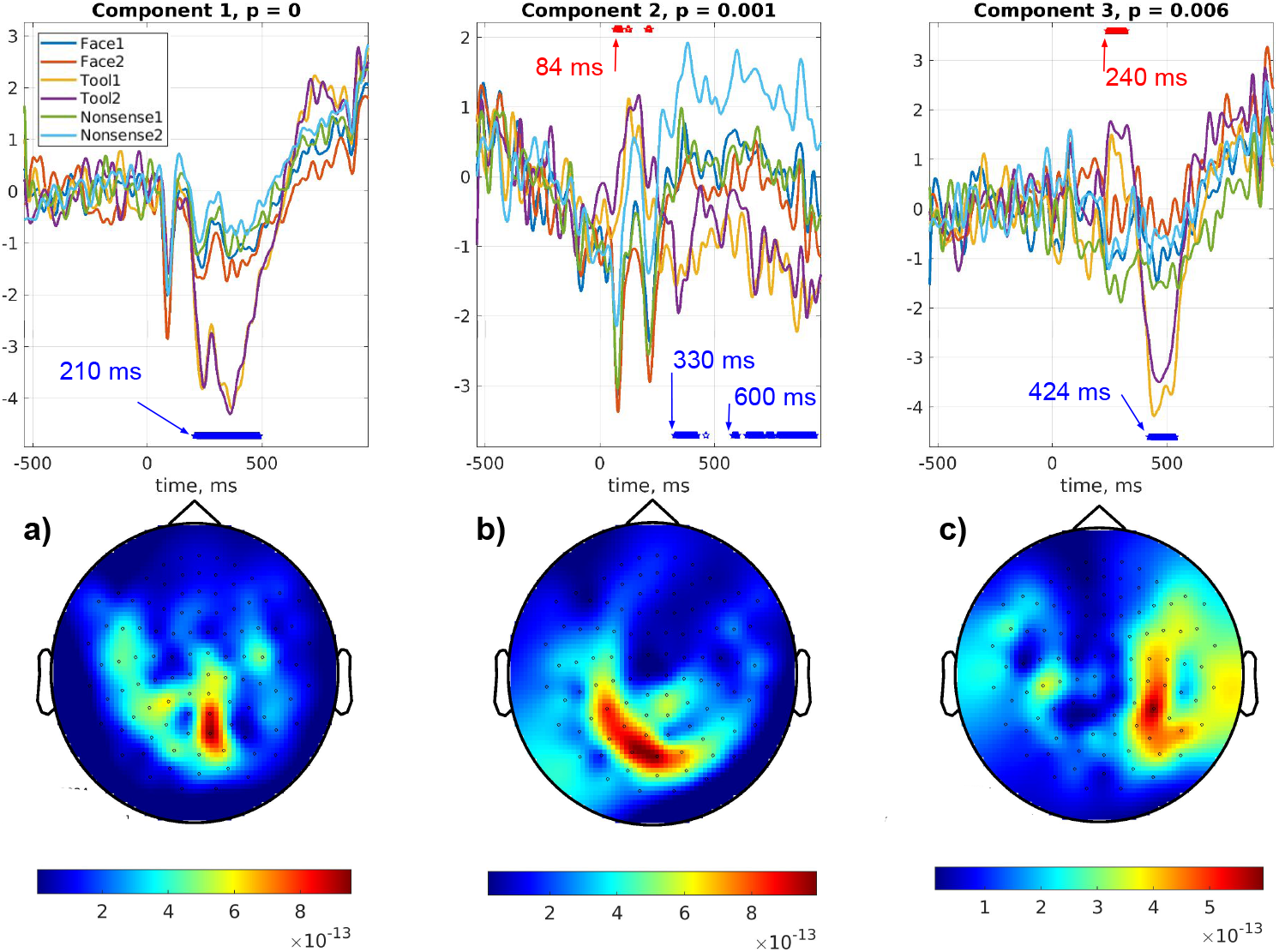
ReDisCA applied to the data from [27] with the tool-specific theoretical RDM, see Figure 12.b.

**Figure 15:**
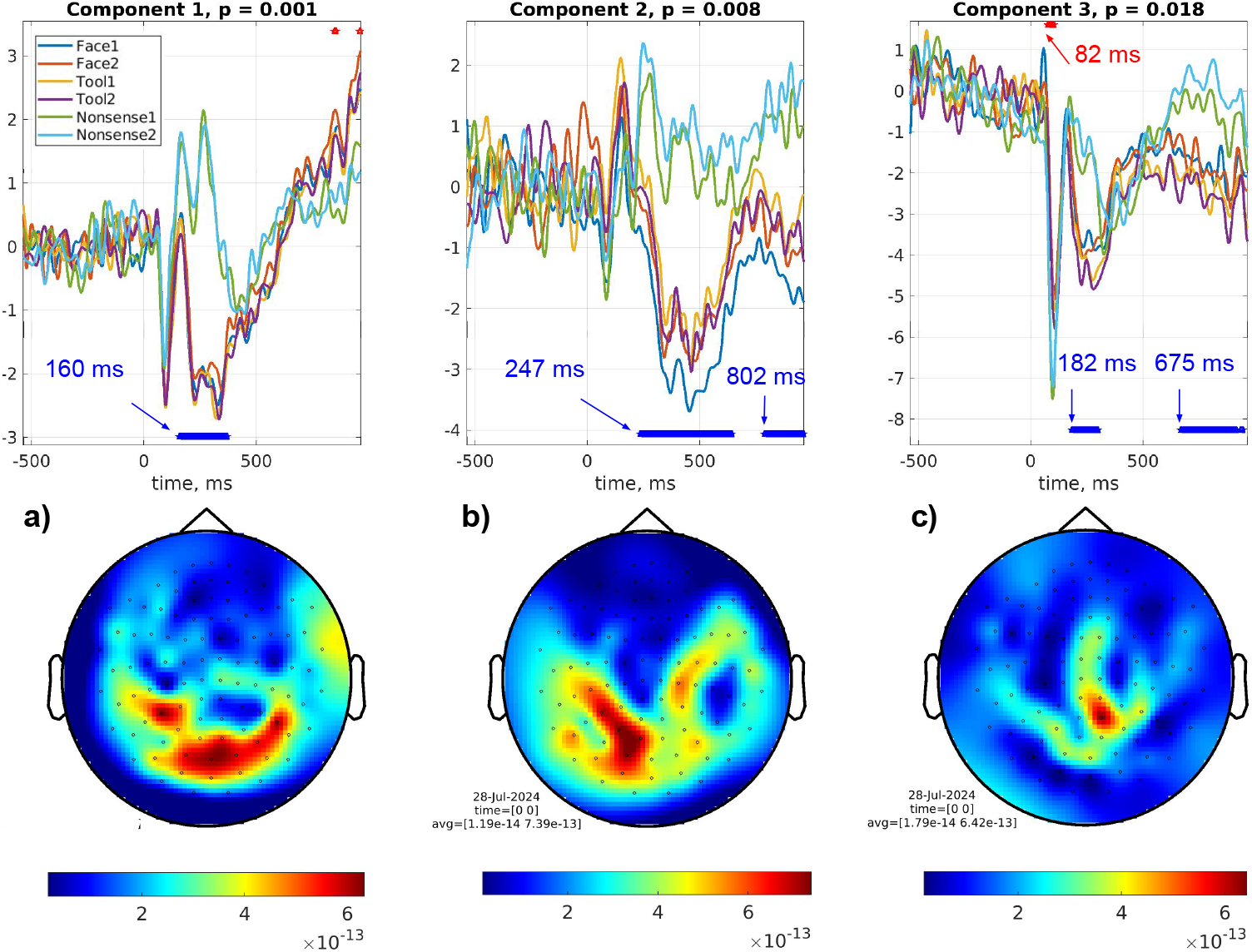
ReDisCA applied to the data from [27] with the RDM contrasting meaningful (“face”,”tool”) vs. meaningless (“nons.”) categories, see Figure 12.c.

The first face category-related ReDisCA component, shown in Figure 13.a exhibits the early differential response to faces starting at 65 ms, see [55] for a review of possible reasons. The response curve has a prominent peak at 160 ms. This peak differentiates the neural response to face stimuli from the responses to the other categories. The spatial topography shown in the bottom panel of Figure 13.a has maximum over the central and right occipital sensors. This topography likely corresponds to the neuronal sources in the occipital pole, inferior occipital and the right fusiform gyrus. Notably, this ReDisCA component rises again with the significance interval starting at 311 ms. This component aligns well with the blue face-related decoding trace shown in Figure 3 of [27]. The second ReDisCA component, see Figure 13.b, demonstrates prominent activation of the parietal cortex with the first significance interval starting at 218 ms. This topography may reflect sources located in the middle temporal posterior cortex. The last ReDisCA component shows late and sustained processing of face-related information starting at 273 ms by neuronal populations located in the bilateral occipital regions involving some activation of the frontal lobes which may reflect fronto-temporal interactions associated with visual processing [45]. This component augments the first ReDisCA component but does not exhibit early response to the face stimuli.

Figure 14 shows the results of ReDisCA with the tool-specific theoretical RDM from Figure 12.b. ReDisCA component 1 (Figure 14.a) shows a prominent response to the “tool 1” and “tool 2” stimuli that occurs at 210 ms and appears to be localized to the central occipito-parietal region also involving the planar gradientometers located over the left (and compactly over the right) central sulcus areas. Later activation involves the extended parietal and occipital regions (Component 2, Figure 14.b) localized to the mid-occipital and left parietal cortices. Interestingly, ReDisCA’s Component 3 shows activity starting later at around 240 ms and stemming from the right and left sensory-motor cortices and sources in the temporal lobe with the prevalence of the right side in contrast to Component 1. These observations align well with those in [27] where the authors also observed responses at similar latency with the left sensory-motor cortex (ReDisCA’s component 1) leading the right one (ReDisCA’s component 3), see the tools-related curves in Figure 3 of [27] for the IPS, VPM and the MTp cortices.

Next, we explore ReDisCA’s components contrasting the perception of the meaningful (“face” and “tool”) and the meaningless (“nons.”) visual stimuli, see Figure 12.c for the corresponding theoretical RDM. In Figure 15.a and Figure 15.b we can see the two components that can be assigned to the ventral and dorsal visual pathways correspondingly. The activity response to the meaningful stimuli occurred as early as 160 ms in the occipito-parietal region (Component 1) followed by the prominent response along the dorsal visual pathway (Component 2). Early (182 ms) and short-lasting as well as the late (675 ms) differential response between the meaningful and the meaningless stimuli gets isolated into a single component with a focal mid-parietal topography, see Component 3 in Figure 15.c.

In the final example we used ReDisCA to impose specific geometric relationships between the resultant components. To do so we used the non-binary RDM shown in Figure 16.a which forces ReDisCA to look for the components whose activation is similar within each of the three categories (faces, tools, and visually meaningless images). Simultaneously, it imposes geometric relations on the activation of the sought components, requiring that the distance between each meaningful category and the meaningless one is less than the distance between the two meaningful categories, see Figure 16.b. Since ReDisCA currently searches for one-dimensional components, the responses to the two meaningful stimuli (faces and tools) will be positioned on the opposite sides relative to the responses to the nonsensical visual stimuli.

**Figure 16:**
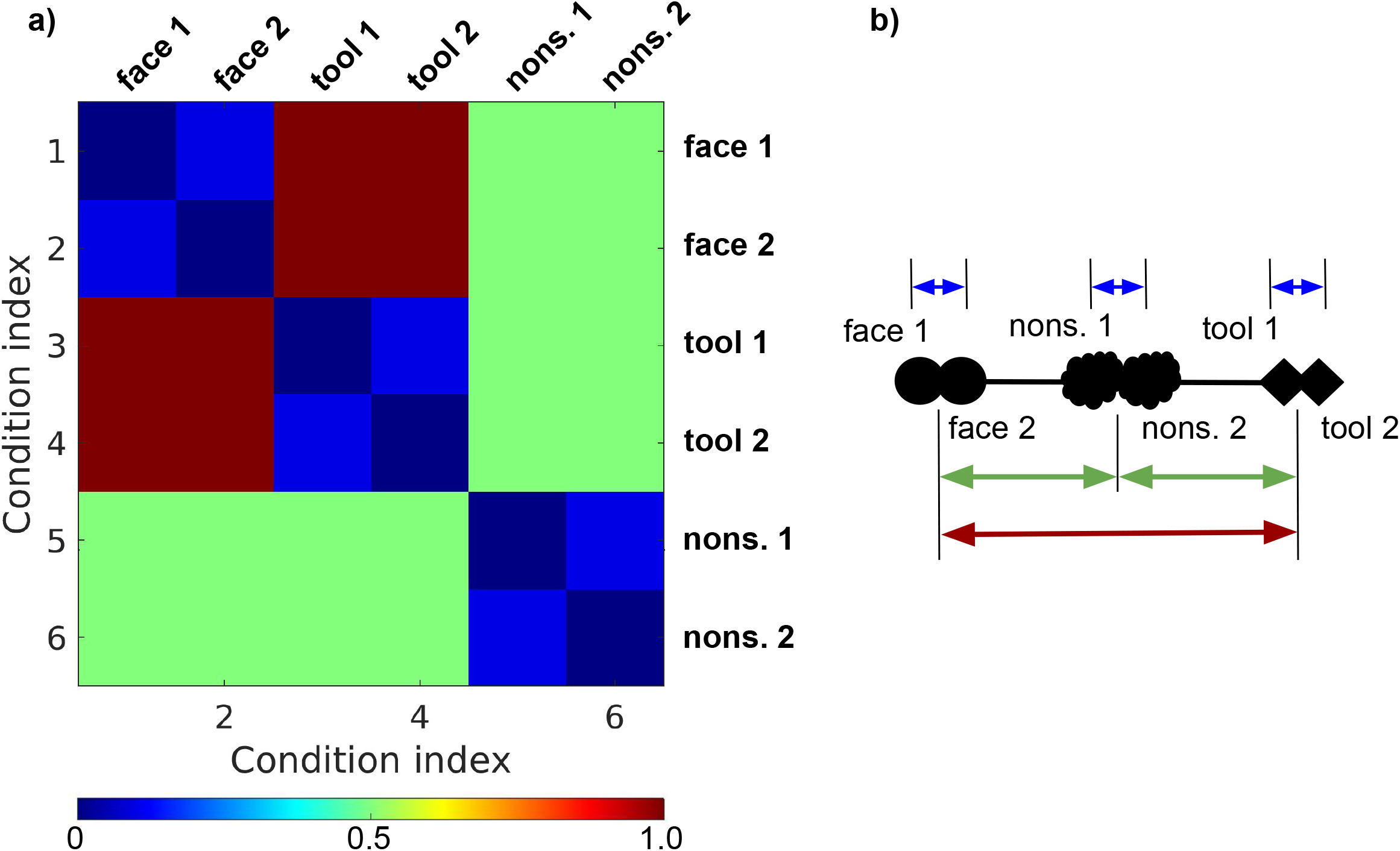
a) The non-binary theoretical RDM matrix used to impose the specific geometric relationship on the component time series. b) This matrix requires that the responses to “face” and “tool” categories are maximally separated while both are equally distanced from the response to the nonsensical images. The distances in a) and b) are identically color-coded.

The three components with the lowest *p*-values are shown in Figure 17 in the order they were returned by ReDisCA.

**Figure 17:**
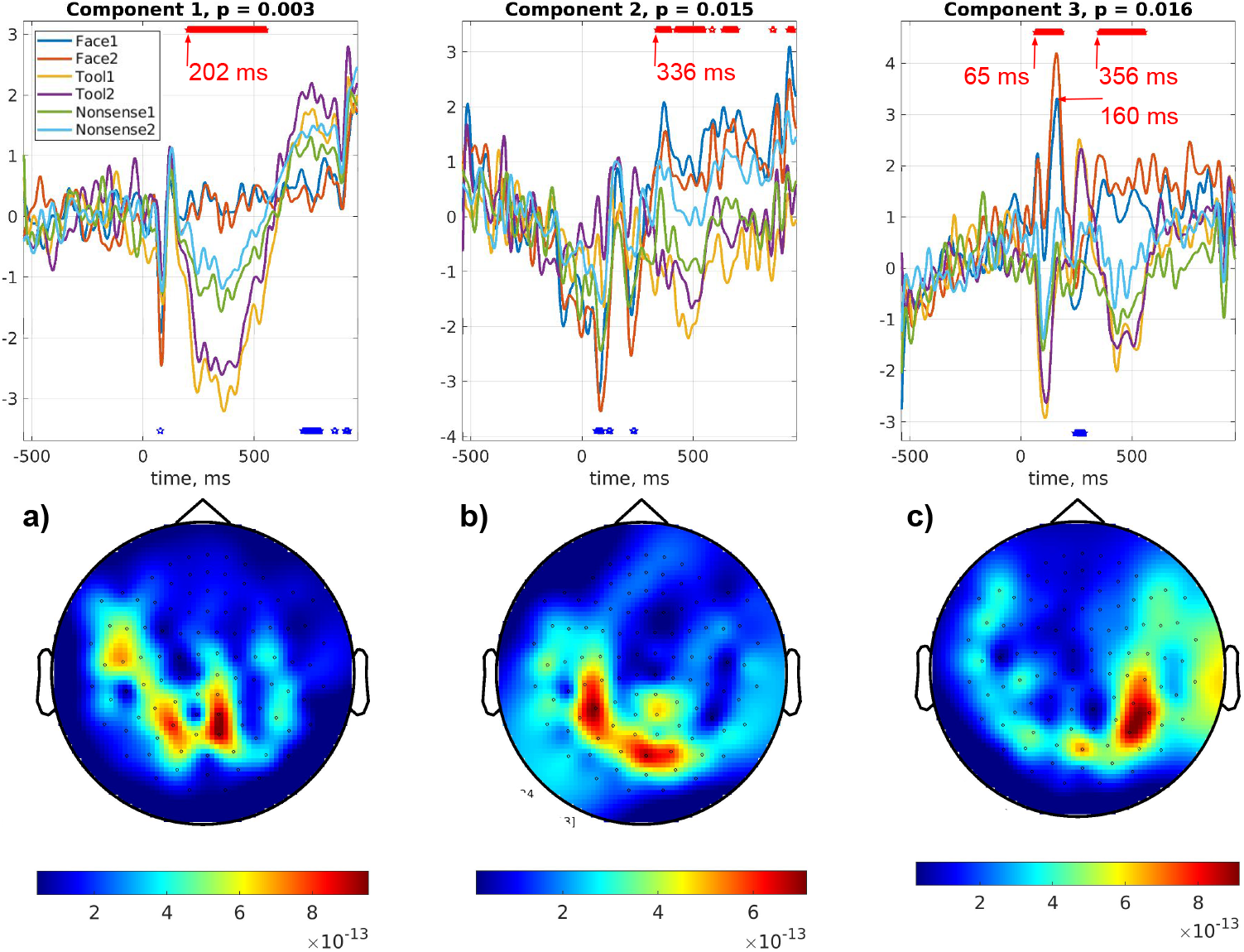
Representational dissimilarity components in the dataset by [27] obtained using non-binary theoretical RDM shown in Figure 16.

From 17.a we can observe the rise of the response magnitude in the “tools” condition that is significantly different from the response in the “faces” condition (that remains nearly flat over this interval) as shown by the asterisks on the top of the plot. The significant difference starts at 202 ms which nearly exactly matches the results reported in Figure 3 of [27]. As evident from the topography, the observed response is produced by dominantly left parietal sources and possibly sources from the left sensory-motor cortex. Component 2 shows the development of this response in the visual areas of the cortex which again aligns well with the late (around 260-350 ms) activity of the fusiform gyrus (FG) in response to the images of “tools”. The last component reflects face-specific activity that peaks early at around 160 ms, then mixes with the activity in the other subcategories and then in the second window starting at 356 ms becomes prominently distinct from the response in the other categories.

We have also performed MUSIC scan in order to highlight the cortical areas whose source topographies fall into the representational dissimilarity subspace identified by ReDisCA. To do so we have utilized the forward model built using the individual MRI as described in [27] and applied equation (14) to compute the subspace correlation 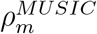 between the pair of topographies corresponding to a freely oriented dipole at each *m*-th cortical location and the representational dissimilarity subspace identified by ReDisCA spanned by the topographies shown in Figure 17.

In Figure 18 we can observe involvement of the dominantly right fusiform gyrus, right insula, left intraparietal sulcus and anterior central gyrus as cortical regions demonstrating the requested geometric relationships imposed by the RDM in Figure 16.a In these regions the response to faces and tools appears on the opposite sides w.r.t. their response to nonsense stimuli as schematized in Figure 16.b and observed in the top panels of Figure 17.

**Figure 18:**
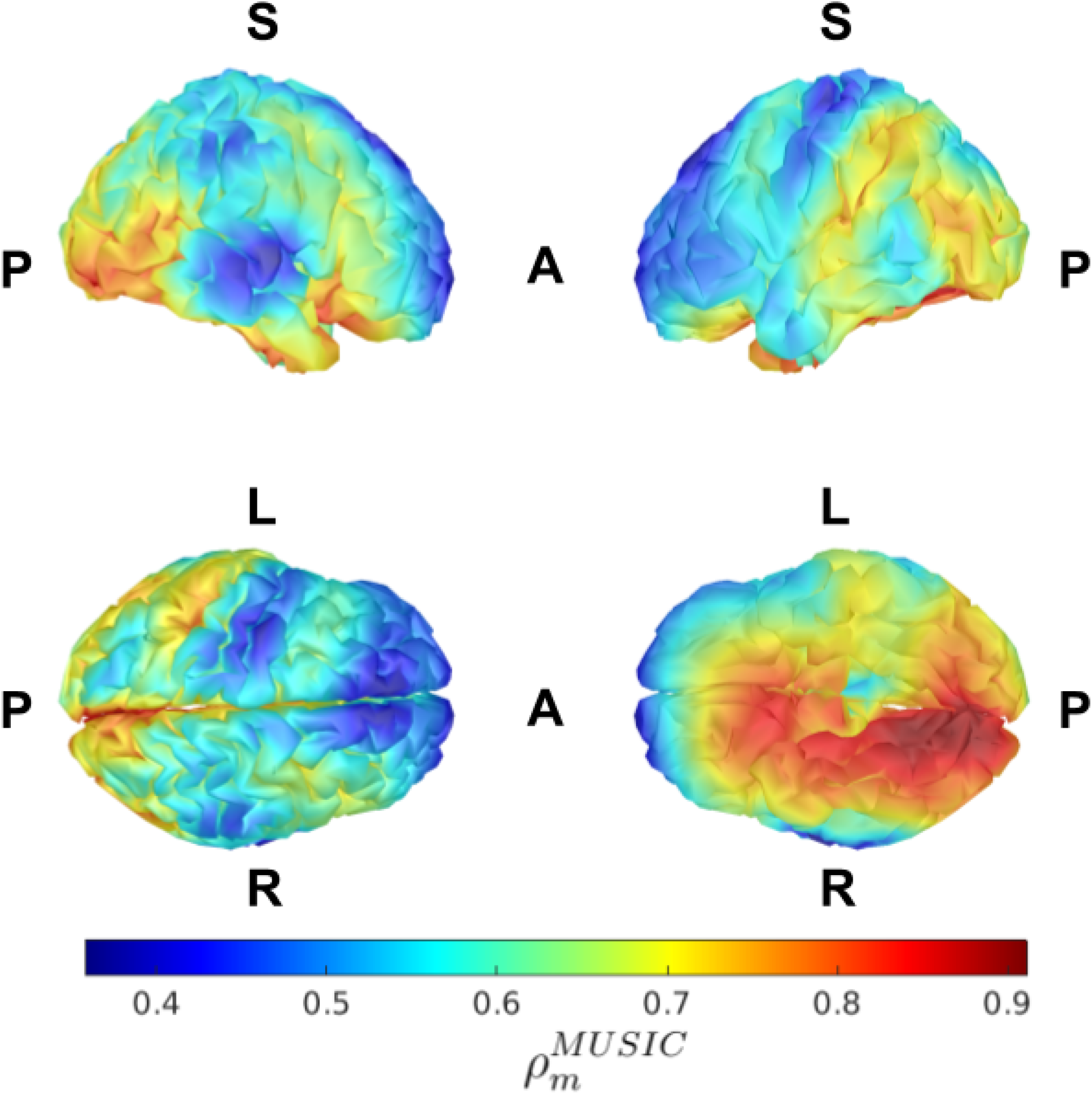
Cortical map reflecting the first principal angle between the dissimilarity subspace spanned by the topographies from Figure 17 and the subspaces spanned by the topographies of a freely oriented dipole at each cortical location, see equation (14). This highlights the potential contribution of the cortical sources to the multivariate activity pertinent to the user specified RDM. (14). Letters A,P,S,L,R - anterior, posterior, superior, left, right define the view angle.

Overall, application of ReDisCA to the MEG dataset exploring the neural processes underlying visual categorization of faces and tools appears to support the main result of the original study [27] and confirms that while perception of faces is dominantly accompanied by the activity in the ventral visual stream, the images of tools elicit activity along the dorsal visual path and also result in the activation of hand related sensory-motor areas associated with the use of the tools.

## 5. Discussion

Conceptually, ReDisCA extends the family of the previously developed methods for decomposing multichannel EEG and MEG data and extracting the spatial-temporal components with desired properties. One popular example is the Spatial Spectral Decomposition (SSD) technique [41]. SSD seeks to maximize the power at a frequency band of interest while simultaneously minimizing it within the side bands. For each specific triplet of frequency bands (one central and two flanking) the SSD results in a set of spatial components whose time series exhibit the frequency band selective property. In particular, the timecourses of the SSD-derived spatial components are characterized by the maximal ratio of power within the central band to that within the two flanking sub-bands which justifies the use of SSD for extracting rhythmic components from multichannel EEG or MEG time series. Another popular method is the Source Power Comodulation, or SPoC [9]. SPoC aims to find spatial filters and patterns by using in the decomposition process a target behavioral variable to emphasize the components whose instantaneous power profile correlates with the target variable. Typically SPoC targets rhythmic activity and extracts its sources whose envelope aligns well with the target variable. Notably, neither SSD nor SPoC or their early fore-runner ICA require forward and inverse modeling machinery to extract target component time series. Theoretically speaking, the design of either of these methods does not require that the obtained components correspond to a specific neuronal source with a well-defined location. In practice, however, the obtained patterns that pertain to individual components exhibit remarkable alignment with the appropriate dipolar electromagnetic model and allow for pinpointing a specific cortical location as a source of activity with the desired property.

In its current version ReDisCA operates with evoked response data recorded over several experimental conditions and aims at finding spatial components (each potentially corresponding to a distinct neuronal source) whose activation time series exhibit the expected representational structure. We have started with the Euclidean norm as a distance between component activation time series. This continuous measure of dissimilarity not only allows for an analytic solution of the subsequent optimization problem but appears superior [58, 15] to the simplest forms of more intricate classification accuracy-based distances. Based on this and using the standard notion of spatial filtering of multichannel EEG or MEG data we devised ReDisCA as the approach for dissecting the evoked responses into a set of spatial components with activation time series that adhere to the user-supplied target (or theoretical) representational dissimilarity structure.

To achieve this we have formulated the specific constrained optimization problem (5). Its solution gives us the spatial filters whose application to the multichannel evoked response yields component activation timeseires with the expected target representational structure encoded by the prespecified target RDM. These time series combined with the spatial patterns corresponding to each of the spatial filters [17, 48, 23] give us spatial decomposition of the multichannel evoked responses observed over the range of experimental conditions. The goodness of fit is given by the corresponding generalized eigenvalue.

In contrast to the source space RSA [27] ReDisCA does not require inverse modeling and does not perform an explicit and exhaustive scan over brain sources in the attempt to pinpoint those whose empirical RDM is close enough to the target RDM. Conceptually the role of the inverse modeling in ReDisCA is similar to that in the multivariate pattern analysis of EEG or MEG data, see for example [7]. While the inverse modeling can be applied to the patterns [17, 24] obtained from the regression weights it is not the essential step in the analysis.

In simulations to compare ReDisCA to the source space RSA we fitted an electromagnetic model to the signal subspace spanned by the topographies of the statistically significant ReDisCA components, see equation (14) and matched the obtained locations against those of the simulated sources. This allowed us to rigorously compare ReDisCA with the explicit inverse modeling-driven RSA. ReDisCA has successfully identified the signal subspace corresponding to the simulated sources and provided higher ROC AUC scores as compared to the source space RSA based on the exhaustive scan. Note that in our implementation of the source space RSA we analyzed each cortical source separately, used MNE and LCMV beam-former as inverse solvers and employed the L2-norm as the dissimilarity measure instead of sLORETA and LDA classifier used in [27]. However, when matching these two techniques against each other one has to keep in mind that ReDisCA uses less information and does not require inverse modeling which makes it applicable to a broader collection of datasets and modalities. For example, ReDisCA may be an interesting option to employ for the analysis of ECoG data where forward and inverse modeling are less straightforward than it is with non-invasive EEG or MEG data. Also, ReDisCA is directly applicable to the averaged evoked responses time series and does not require the presence of single-trial data.

In the final example of real-data analysis we have visualized the representational dissimilarity subspace discovered by ReDisCA. To this end we have performed MUSIC scan and visualized the cosine of the first principal angle between the dissimilarity subspace and the subspaces spanned by the topographies of a freely oriented dipole at each cortical location. This highlights the potential contribution of the cortical sources to the multivariate activity pertinent to the user specified RDM. More sophisticated analysis of the ReDisCA discovered subspace can be performed using a variety of EEG and MEG inverse problem solving approaches [14] including the advanced hierarchical Bayesian techniques aimed at recovery of the underlying neuronal current source density e.g. [33, 50] or parametric approaches to discover discrete sources [35] and their associated time series. Doing so one has to take into account the importance of accurately modeling individual subject heads and tissue conductivity profiles [59] to improve the source localization accuracy. At the same time, ReDisCA, a spatial decomposition method, as opposed to the source-space RSA, does not require the inverse modeling and can be used to analyze the datasets with no detailed anatomic information, see Section 4.2.1. Although at the expense of potentially important details, in such a setting ReDisCA can be useful in non-invasive explorations of the representational drift phenomenon [11] and in the analysis of psychiatric conditions e.g. [53].

Any RSA implementation benefits from the increase in the number of experimental conditions. Although we demonstrated that ReDisCA outperformed conventional RSA approaches for all tested counts of conditions (3,4,5,6), the practically useful results can be obtained from datasets with no less than four conditions. MNE inverse solver appeared to be a better option as compared to the LCMV beamformer for the number of conditions less than 6. ReDisCA followed by the source localization procedure yielded almost a 2-fold reduction in the mean source localization error as compared to the exhaustive scanning source space RSA. The approach reported in [27] where voxel time series obtained with sLORETA are combined into ROIs followed by dimension reduction and LDA classification. This combination may potentially improve the accuracy of the exhaustive scan-based RSA however, requires significantly more computations and in contrast to ReDisCA is not directly applicable to the averaged evoked responses time series.

We demonstrate that the optimization problem in ReDisCA is mathematically analogous to that of SPoC, but it utilizes modified input data matrices. These matrices are calculated as the difference between the evoked responses recorded for each pair of conditions. In this context, the entries of the theoretical RDM formally take on the role of the behavioral variable, as shown in Table 1.

In the future, ReDisCA can potentially adopt cross-validated Euclidean distances [58]. To cope with the inherent asymmetry one can use a symmetric version of it as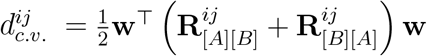, where 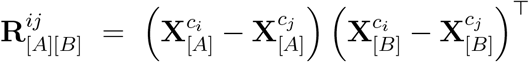 and 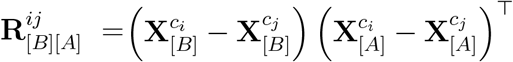 with 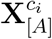 representing averaged responses corresponding to condition *c*_*i*_ and obtained using base data partition A and 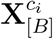 - using validation partition B. Investigation of the utility of this cross-validated distance in the ReDisCA setting is a separate topic that needs to be addressed by future studies.

Feature-weighted RSA is a modification of RSA that allows to better align the observed and the theoretical RDMs [21]. This is achieved by mixing the RDMs that pertain to the individual features, e.g. voxels comprising an ROI, with optimal weights maximizing the similarity of this mixed RDM and its theoretical target. In application to EEG and MEG data the role of individual features is played by individual channels and therefore the adaptively tuned weights may be considered as some sort of spatial filtering. At the same time, since the weighted RDM matrix entries appear in the form of non-linearly transformed channel data (squared distances, functions of classification accuracy, etc.) the coefficients of such a mixing can not be transformed into electromagnetically meaningful patterns and are therefore difficult to interpret [13].

In contrast, ReDisCA by design seeks for a linear spatial filter applied directly to the channel data to isolate a source with the desired representational pattern. Because of this, the weights vector can be easily converted to source topography, or pattern [24, 17], which takes into account the spatial correlation structure present in the data. This guarantees that the discovered spatial patterns can be used for implementing a rigorous source localization procedure based on the notion of signal subspace as described for example in [40]. This property positively distinguishes ReDisCA from the sensor space RSA approach described in [37] where the authors using high-density EEG were able to highlight the detailed spatial and temporal patterns of responses underlying various aspects of meaningfulness in the presented visual stimuli. They achieved this using newly proposed differentiation analysis (DA) and computed temporal and spatial profiles of the category differentiation index (CDI) that can be used to make judgments about time windows and approximate spatial locations of the pivotal sources. However, due to their empirical nature, the obtained spatial CDI maps can not be rigorously fitted with an electromagnetic model and therefore are only tangentially related to the underlying distribution of cortical sources whose activity exhibits the desired representational properties.

ReDisCA is formulated to search for a single spatial vector, as indicated by Equation (5), and evaluates the similarity of activation in the temporal dimension. This places specific constraints on the size of the temporal window *T* to ensure ReDisCA’s proper functioning. It’s crucial to recognize that the size of this window is contingent upon the frequency content of the investigated evoked response. Faster-changing signals permit more information (pertaining to similarity/dissimilarity) within shorter time intervals.

Additionally, the window size is influenced by the number of sensors employed. In theory, larger sensor arrays necessitate longer windows to ensure the correlation matrices are full rank. Note, however, that typically, EEG and MEG data are projected into smaller subspaces with dimensions 10-100. All subsequent operations are conducted within this reduced dimensional space to guarantee the full rank of the resulting matrices. An alternative or additional approach involves employing the Tikhonov regularization strategy when computing the inverse of the correlation matrix 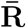. It’s evident that reducing the subspace size and increasing the regularization parameter, in addition to enhancing stability, adversely impacts ReDisCA’s spatial resolution. At the same time, as illustrated with the last example described in Section 4.2.2 ReDisCA can be applied to the entire response duration window and automatically provides the time-intervals where the sought representational dissimilarity structure arises. Guidelines detailing these trade-offs must be developed in the future, drawing upon both simulations and theoretical analyses followed by verification with real data.

The approaches such as SSD, SPoC and now ReDisCA are designed to be applied to a single subject. When the conclusion needs to be drawn from a cohort of subjects, special steps need to be undertaken. For example, in the source space RSA approach reported in [27] pooling across subjects is performed using the standard in the MEG approach based on warping individual cortex geometries onto the canonical cortex followed by calculation of the group level statistics. Given the superiority of ReDisCA demonstrated here as compared to the explicit scan in detecting sources with specified RDMs, see Figures 4 and 5, the source space estimates can be obtained by applying the individual inverse modeling to the patterns discovered by ReDisCA followed by warping these individual results onto the canonical brain. It is potentially possible to consider development of group-level approaches, conceptually similar to those reported in [28, 20] and driven by ReDisCA functional (5).

ReDisCA’s applications extend beyond EEG and MEG data analysis to potentially include the ROI-based examination of fMRI data. Interpreting the weights that scale individual voxel RDMs, as introduced in [21], poses challenges in converting it into a pattern, as discussed in [17]. Here, the difficulty arises because the weights in [21] are applied to the Representational Dissimilarity Matrices (RDMs) of individual voxels and not to the actual voxel activations. Since the RDM elements are non-linear functions of voxel activation, the weights in [21] can not be converted into within ROI activation patterns highlighting the voxels that contribute most to the target RDM.

Employing ReDisCA within each ROI may resolve this challenge, by applying weights to the activation of individual voxels rather than to their non-linear functions. This would enable the subsequent transformation of ReDisCA weights into patterns [17, 46] which could enhance the spatial resolution of fMRI-based functional mapping by clarifying the delineation of an extended ROI into subregions contributing to the improved alignment with the target RDM.

Moreover, ReDisCA can advance the methodology outlined in [32], where the RSA principle was used to analyze representations emerging in the layers of an artificial neural network. An intriguing possibility arises when ReDisCA-derived weights align with those of a subsequent neuron connected to the analyzed layer, signifying that the individual neuron appears to be tuned to track the representational structure encoded in the target RDM.

## 6. Conclusion

In conclusion, this study introduces and applies the novel technique of Representational Dissimilarity Component Analysis (ReDisCA) to overcome challenges associated with classical Representational Similarity Analysis (RSA), particularly in EEG and MEG data. By estimating spatial-temporal components aligned with a target representational dissimilarity matrix (RDM), ReDisCA produces spatial filters and associated topographies that reveal “representationally relevant” sources. The application of these spatial filters to evoked response time series demonstrates superior performance in terms of source localization accuracy compared to conventional source space RSA, with significantly fewer computations. Our approach does not require forward modeling and any geometric information but if these are available the obtained components permit rigorous fits with the electromagnetic model by means of a plethora of methods for solving the inverse problem. The methodology is validated through realistic simulations and applied to a real EEG and an MEG dataset, showcasing its potential for discovering representational structures without relying on the inverse modeling. In the discussion, we highlighted the limitations of ReDisCA, outlined potential directions for its further development, and identified novel applications.

## 7. Funding

The article was prepared within the framework of the Basic Research Program at HSE University.

## 8. Conflict of interest statement

The authors declare that they have no conflicts of interest regarding the publication of this article.

## Notes

https://osf.io/pfde9

### Competing Interest Statement

The authors have declared no competing interest.

### Summary of Updates

added analysis of an additional dataset from (Kozunov et al. 2018).

